# Reprograming human fibroblasts into Sertoli cells: a tool for personalized medicine

**DOI:** 10.1101/2022.08.25.505300

**Authors:** Abhinav Parivesh, Emmanuèle Délot, Alejandra Reyes, Janelle Ryan, Surajit Bhattacharya, Vincent Harley, Eric Vilain

## Abstract

Disorders/Differences of Sex Development (DSD) are congenital conditions in which the development of chromosomal, gonadal, or anatomical sex is atypical. With overlapping phenotypes and multiple genes involved, poor diagnostic yields are achieved for many of these conditions. The current DSD diagnostic regimen can be augmented by investigating transcriptome/proteome in vivo, but it is hampered by the unavailability of affected gonadal tissue at the relevant developmental stage. We try to mitigate this limitation by reprogramming readily available skin tissue-derived dermal fibroblasts into Sertoli cells (SC), which could then be deployed for different diagnostic strategies. SCs form the target cell type of choice because they act like an organizing center of embryonic gonadal development and many DSD arise when these developmental processes go awry.

We employed a computational predictive algorithm for cell conversions called Mogrify to predict the transcription factors (TFs) required for direct reprogramming of human dermal fibroblasts into SCs. We established trans-differentiation culture conditions where stable transgenic expression of these TFs was achieved in 46, XY adult dermal fibroblasts using lentiviral vectors. The resulting Sertoli like cells (SLCs) were validated for SC phenotype using several approaches. SLCs exhibited Sertoli-like morphological and cellular properties as revealed by morphometry and xCelligence cell behavior assays. They also showed Sertoli-specific expression of molecular markers such as SOX9, PTGDS, BMP4, or DMRT1 as revealed by IF imaging, RNAseq and qPCR. The SLC transcriptome shared about two thirds of its differentially expressed genes with a human adult SC transcriptome and expressed markers typical of embryonic SCs. Notably, SLCs lacked expression of markers of other gonadal cell types such as Leydig, germ, peritubular myoid or granulosa cells.

The trans-differentiation method was applied to a variety of commercially available 46, XY fibroblasts derived from patients with DSD and to a 46, XX cell line. The DSD SLCs displayed altered levels of trans-differentiation in comparison to normal 46, XY-derived SLCs, thus showcasing the robustness of this new trans-differentiation model.

## Introduction

DSD are defined as “congenital conditions in which development of chromosomal, gonadal, or anatomical sex is atypical”(Hughes et al., 2006; P. A. Lee et al., 2006). They may present in isolation or as part of complex multiorgan developmental syndromes (Baetens et al., 2016; E. C. Délot et al., 2017; Hutson et al., 2014). DSD diagnosis combines several approaches including anatomic description and endocrine and genetic testing. Genetic testing is required to disambiguate many conditions who present with a similar phenotype but can be associated with the dozens of genes identified so far (E. C. Délot & Vilain, 2021; Parivesh et al., 2019)

Genetic testing for DSD spans from traditional karyotyping to chromosomal microarrays, to next generation sequencing approaches (exome or whole genome sequencing), as well as some more specific approaches for genes refractory to short-read sequencing such as *CYP21A2* (E. C. Délot & Vilain, 2021; León et al., 2019). Pre-clinical technology such as optical genome mapping to detect structural variants (Barseghyan et al., 2018) or long-read sequencing offer the hope of future increased diagnosis yield. However, at this time only 30-50% of patients receive a specific molecular diagnosis in the research setting, and much lower in many clinical settings (Croft et al., 2016; E. C. Délot et al., 2017; E. Délot & Vilain, 2018). The low diagnostic rate may be because new DSD-associated genes are waiting to be discovered or because currently available technology misses several types of variants. However a major limitation is also the difficulty in interpreting the significance of variants.

Transcriptome sequencing from blood, muscle, fibroblasts etc., coupled with genetic testing has been shown to significantly increase diagnostic rates in other rare Mendelian conditions(Cummings et al., 2017; Hamanaka et al., 2019; Kernohan et al., 2017; Kremer et al., 2017). However, procuring a relevant tissue to examine the transcriptome is not possible for conditions where pathogenic mechanisms occur early in development as is the case for most DSD conditions. While a biopsy of embryonic gonad is impossible to obtain, skin biopsies can be readily collected for isolating dermal fibroblasts. If these can be reprogrammed into gonadal cell types *in vitro*, they could be used for transcriptomic analysis to identify and validate variants in a patient-specific model. While the fetal gonadal transcriptome is still poorly defined and highly complex and dynamic (Lecluze et al., 2020), deriving patient-specific cell types that approximate those developing cells should provide useful tools for DSD diagnosis.

As the main organizing center of the typical male gonad, Sertoli cells (SC) are central to the etiology of many DSD (Ross & Capel, 2005; Ryohei Sekido et al., 2004; Svingen & Koopman, 2013). They are therefore the prime target choice to reprogram somatic cell type into for patient-specific transcriptomic diagnostic of DSD. Sertoli cell growth and development in humans happens in three phases: immature SC proliferate in the prenatal and neonatal phase of development; they then remain quiescent for 8-10 years; in the third phase, proliferation restarts in the peri-pubertal age, making the final number of mature, non-proliferative SCs of the adult testis which are characterized by different transcription patterns than immature SCs (Cool & Capel, 2009; Mackay, 2000; McLaren, 2000). Expression of several key genes at the genital ridge of chromosomal males triggers a cascade leading to *SRY* expression and to differentiation of Sertoli cells. For example GATA4-FOG2 interaction induces *SRY* expression (Tevosian et al., 2002). *WT1* activates *NR5A1*/SF1 expression early in the bipotential stage of the gonad, which supports induction of *SRY* (Hanley et al., 1999; Wilhelm & Englert, 2002). In the early, bipotential gonad expression of *SRY* is followed by *SOX9*, which triggers the differentiation of primordial SCs. SCs form the physical framework of seminiferous cords, which provide nutritional and structural support to developing primordial germ cells, peritubular myoid cells, endothelial cells. SCs also drive the differentiation of androgen-secreting Leydig cells, which in turn shapes the male pattern of sexual development (Sharpe et al., 2003; Svingen & Koopman, 2013; Tarulli et al., 2012). Variants in all these genes can lead to DSD in humans (E. C. Délot & Vilain, 2021; Parivesh et al., 2019). For example, mutations, deletions, inversions in *SOX9*, as well as in its upstream promoter and enhancer regions cause campomelic dysplasia, a condition associating skeletal malformations and to sex reversal in XY individuals (Cameron & Sinclair, 1997; Croft et al., 2018; Knower et al., 2003; Sekido & Lovell-Badge, 2013; Wagner et al., 1994).

Transient trans-differentiation of pluripotent stem cells (iPSCs) into Sertoli-like fate was achieved with specialized small molecules or biological factor supplementation (Knarston et al., 2020; Rodríguez Gutiérrez et al., 2018). A transgenic approach was able to produce potentially stable Sertoli-like cells using murine fibroblasts (Buganim et al., 2012) but this hadn’t been reproducible with human cells. A recent report attempted human fibroblasts to Sertoli trans-differentiation using a stable transgenic approach with just two transcription factors: *NR5A1* and *GATA4* (Liang et al., 2019). It is however not clearly specified whether the resulting cells express markers of other gonadal cell types, and specificity of the method is difficult to ascertain.

Here we present a method of transgenic expression of a novel combination of transcription favors (TFs) to produce Sertoli-like cells (SLC), as predicted by the computational predictive network Mogrify (Rackham et al., 2016), using lentiviral vectors and suitable culture conditions. We transduced adult 46, XY human dermal fibroblasts, and validated the approach by looking for the appearance of Sertoli-specific molecular signatures and cellular phenotypes. Further, we applied this method to dermal fibroblasts derived from a variety of genetic backgrounds, including 46, XX as well as 46,XY DSD patients to test the robustness of the method in achieving trans-differentiation to a Sertoli-like phenotype.

## Results

### Adult human 46, XY dermal fibroblast to Sertoli cell trans-differentiation strategy produces Sertoli like cells (SLC) that exhibit a significant change in morphology

We utilized the predictive software Mogrify, to predict the TFs necessary to direct differentiation of fibroblasts into Sertoli cells. Mogrify is a comprehensive atlas of human cell conversions that utilizes gene expression data with regulatory network information from ~300 different cell and tissue FANTOM5 data sets to compute a ranked list of transcription factors that cumulatively exert up to 98% differential regulatory influence on the target cell type in comparison to starting cell type (Rackham et al., 2016). Mogrify predicted that transgenic expression of eight TFs, *FGF2, GATA6, GATA4, MXI1, JUNB, NFYB, NR5A1* and *EBF1,* was required for human dermal fibroblast to Sertoli cell trans-differentiation. It also predicted that just six out of those eight, excluding *NR5A1* and *EBF1,* were enough to achieve a cumulative 95% Sertoli influence on dermal fibroblasts (Figure 1A). *Fgf2* and *Gata4* are differentially expressed between male and female gonads of mice at developmentally important stages, as revealed by RNA-seq analyses (Jameson et al., 2012; Zhao et al., 2018). *NR5A1* is a well-known human DSD gene (Buonocore & Achermann, 2016).

**Figure 1:**
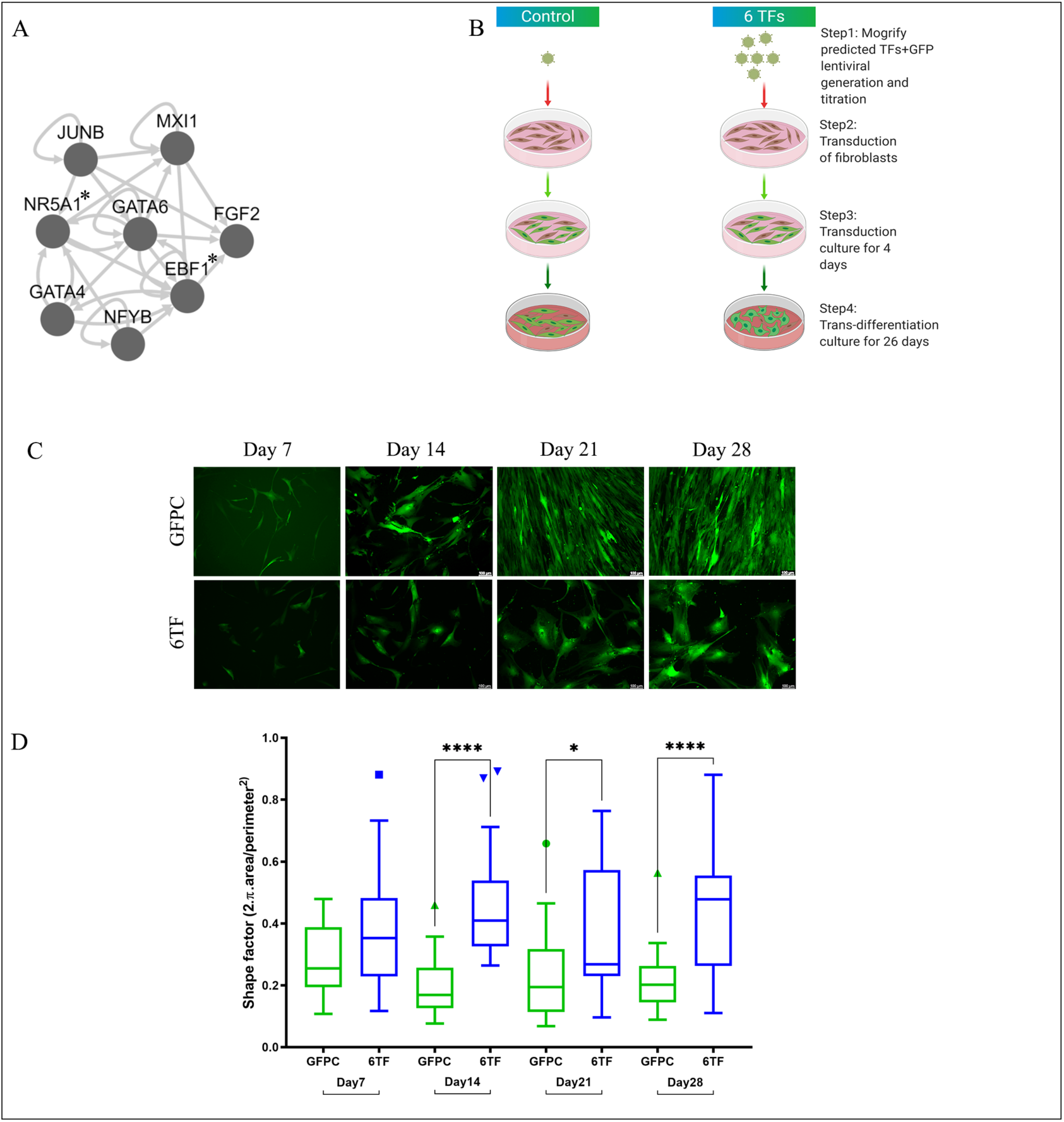
46, XY fibroblast-derived SLCs exhibit significant change in morphology over the 1-month trans-differentiation. (A) Mogrify-predicted transcription factors required for dermal fibroblasts to Sertoli cell trans-differentiation, *=not required to reach 95% coverage of the required genes (B) Outline of the trans-differentiation strategy employed throughout this report. (C) representative images of live cells belonging to the indicated groups expressing GFP across the I-month trans-differentiation culture of 46, XY dermal fibroblasts. (D) Morphometric analysis of cells in (C) showing shape factor quantification represented as Tukey’s box plot with outlier values at either ends of the box (matching colored shapes), drawn from an average of 3 biological replicates (N), counting ~50-60 cells (n) within each group. * represent p values calculated from Mann-Whitney statistical tests conducted between the indicated groups. p > 0.05 (non significant, omitted from the graph), p ≤ 0.05 (*), p ≤ 0.01 (**), p≤ 0.001 (***), p ≤ 0.0001 (****)

In a pilot study, we tested a pilot strategy for trans-differentiation of 46, XY normal adult human dermal fibroblasts using 16 different conditions (Figure 1-figure supplement 1A). stably expressing 8 or 6 TFs (and GFP) at different multiplicity of infections with different transduction and trans-differentiation media. The resulting cells were imaged and harvested to measure expression of Sertoli markers by qPCR. Considering optimal transduction efficiency, cell survival/proliferation, morphological appearance, and expression of Sertoli markers *SOX9*, *PGDS* and *BMP4*, conditions 6 and 5 were chosen to be pursued (and will be called “6TF” and “8TF” respectively below) (Figure 1B).

One of the defining features of cells is their morphology. Of the two cell types relevant to this study, fibroblasts exhibit an elongated spindle-shaped morphology, while Sertoli cells are known to assume squamous epithelial morphology. Sertoli cells grown in adherent cultures have been reported to appear large, have a polygonal shape and exhibit long cytoplasmic extensions or filamentous processes of irregular lengths radiating out of the cell body, along with irregularly shaped nucleus (Chui et al., 2011; Lakpour et al., 2017; Shlush et al., 2017; Wen, Yuan, Sun, Niu, Wang, Fu, Zhou, Yao, Wang, Li, & He, 2017). Cells were imaged weekly under green fluorescence for the duration of the one-month culture (Figure 1C). The size and shape of cells were investigated by morphometric analysis of cell images with recording of the surface area and perimeter of the individual cells.

Shape factor measurement revealed that while there was no significant difference in the shape between GFP transduced control and 6TF, 8TF SLCs in the first week, the shape of 6TF and 8TF SLCs had diverged significantly from GFP control starting at the 2^nd^ week (Figure 1D, Figure1-figure supplement 1D). While the GFP control cells remained elongated in shape, characteristic of fibroblasts, 8TF cells started to assume more polygonal appearance with elongated and irregular cytoplasmic processes, typical of Sertoli cells. These irregular processes were even more accentuated in 6TF cells, as observed visually and quantified by a lower shape value in 6TF SLCs than the more circular or polygonal 8TF SLCs (Figure 1-figure supplement 1D). Additionally, the GFP control and 6TF, 8TF cells appeared to be similar in size in the first week but the experimental cells started flattening, spreading and growing in size in comparison to GFP control beginning in the second week and this difference became even more enhanced with each passing week (Figure 1C, Figure 1-figure supplement 1D). All the cells, including GFP controls exhibited significant growth in size at the end of 1 month in comparison to when they started (Figure 1-figure supplement 1D). Together, these morphometric evaluations showed that the 1 month of trans-differentiation culture gradually pushed fibroblasts to change in size and shape and to start exhibiting some morphological features associated with Sertoli cells.

### SLCs exhibit Sertoli molecular signatures

To investigate the Sertoli features of transdifferentiated cells further we compared their transcriptome with that of a published adult Sertoli cell (aSC) transcriptome (Wen, Yuan, Sun, Niu, Wang, Fu, Zhou, Yao, Wang, Li, He, et al., 2017), and curated lists of Sertoli marker genes (Table S1).

Principal component analysis (PCA) of RNA seq transcriptomes revealed that replicates of 6TF SLC cluster away from GFPC (Figure 2A), while 8TF did not (Figure 2-figure supplement 1A). Replicates of each category of samples grouped together and away from samples of other category (Figure 5-figure supplement 1A).

**Figure 2:**
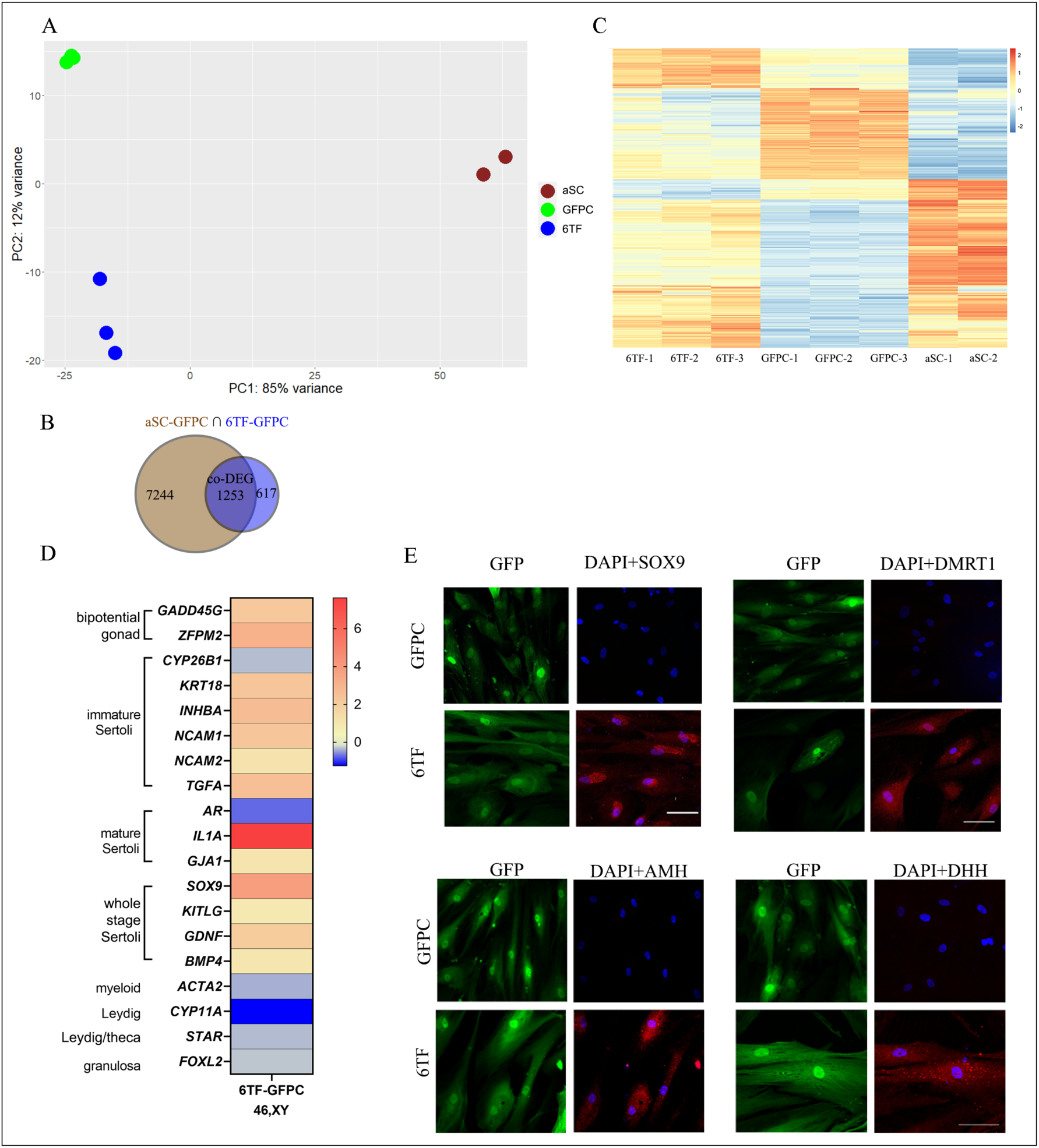
46,XY SLCs show Sertoli marker gene expression: Whole-genome transcriptional profiling of adult Human dermal fibroblasts (HDFa)-derived SLCs. (A) Principal Component Analysis of the transcriptomes of triplicate samples for GFP control (in green), 6TF (in blue) and a published (Wen, et al., 2017) adult Sertoli cell (in brown; note that only 2 replicates were used in that publication). (B) Venn diagram displaying differentially expressed genes (DEGs, fold change>1.5, p adjusted <0.05) between GFPC (N=3) vs adult Sertoli cells (aSC) (N=2) and GFPC (N=3) vs 6TF SLCs (N=3). Intersection part shows co-differentially expressed genes (co-DEGs). (C) Expression of the 1253 co-DEGs in 6TF, GFPC and aSC replicates. (D) Heat map showing 6TF vs GFPC log2 fold change for indicated gonadal marker genes measured by RNAseq (n=3). (E) Representative immuno-fluorescence images showing subcellular expression of SOX9, DMRTl, AMH and DHH in the 6TF SLCs and GFP controls

To normalize potential differences owing to their different origins, all three groups (6TF, 8TF and aSC) were first compared with 46,XY-derived GFP control (GFPC) to determine the set of differentially expressed genes (DEGs). Of the 1870 genes differentially expressed between 6TF and control GFPC, 2/3 were in common with the aSC DEGs (Figure 2B). In 8TF cells, this percentage was almost 3/4 (502/681 genes) (Figure 2-figure supplement 1B). Such high percentages of co-differentially expressed genes between the transcriptomes of SLCs and adult Sertoli cells indicated that our experimental SLCs had significant Sertoli-like gene expression patterns.

Gene ontology (GO) analysis of the co-DEGs determined above identified several terms expected to be associated with Sertoli cell as organizing center of male reproductive system structure and seminiferous tubule development (Figure 2 figure supplement 1D). The top twenty GO biological processes list included *developmental process*, *organ development*, *urinogenital system development*, *tube morphogenesis*, *external encapsulating structure organization*, *ECM structure and matrix organization*, *tube development*, with *reproductive system development*, and *reproductive structure development* further down the list. It also showed terms for positive and negative regulation of lipid biosynthetic and metabolic processes, congruent with the known lipid metabolism and lipid droplet formation in Sertoli cells (Bucay et al., 2008; Chui et al., 2011).

Next, we compared the gene expression profile of the 1253 co-DEG in 6TF SLC, GFPC and aSC in a heatmap (Figure 2C). The 6TF SLC replicates appear quite different from GFPC and notably, resemble aSC expression to a large extent. In contrast, a similar heatmap looking at expression of the 502 8TF-GFPC co-DEG genes revealed that 8TF samples did not show as high resemblance with that of aSC (Figure 2-figure supplement 1C). Thus 6TF might be a better condition for SLC trans-differentiation than 8TF.

In order to investigate the molecular phenotype, a gene list of immature, mature, and all-stage Sertoli markers was drawn up from literature (Table S1). The list also includes markers for non-Sertoli male gonadal cell types such as Leydig, myeloid, and germ cells, as well as for granulosa cells, the female homologue of Sertoli cells.

6TF SLC exhibited an increase in expression of bipotential gonad markers *GADD45G*, *ZFPM;* immature Sertoli markers *KRT18, INHBA, NACAM1/2, TGFA;* mature Sertoli markers *IL1A, CX43/GJA1*; and all-stage-Sertoli markers *SOX9, KITLG, GDNF, BMP4* over GFPC (Figure 2D, Figure 2-figure supplement 1E).

In contrast, myeloid (*ACTA2)* and granulosa (*FOXL2)* cell markers did not show an increase; Leydig cell markers *CYP11A1* and *STAR* decreased (Figure 2D). (Expression of germ cell marker *DDX4* was undetected; data not shown). This indicates that the six transcription factors *GATA4, GATA6, MXI1, JUNB, NFYB* and *FGF2* were sufficient to push dermal fibroblasts towards a Sertoli fate with predominant expression of immature and all-stage markers.

8TF SLC exhibited a significant increase in the expression of bipotential gonad marker *GADD45G*, immature Sertoli markers *CYP26B1, PTGDS, KRT18,* and *NCAM1/2,* and all-stage Sertoli markers *SOX9* and *BMP4* over GFPC (Figure 2-figure supplement 1E). 8TF SLC also exhibited a similar pattern for mature markers (*CTSL, IL1A* and *CX43/GJA1)* with lower fold increase than for immature markers. While 8TF SLCs also didn’t show an increase in expression of myeloid marker *ACTA2* or granulosa marker *FOXL2*, but they did exhibit an increase in expression of Leydig markers *HSD3B2*, *CYP11A* and *STAR* (Figure 2-figure supplement 1E). Unexpectedly high expression of Leydig markers in 8TF in comparison to 6TF suggests that 6TF is a better condition for producing SLCs as 8TF additionally produce Leydig cell characteristics. We will therefore focus on 6TF in the rest of the report (with results for 8TF shown mostly in supplementary figures).

Expression of some markers was further validated by qPCR, which confirmed the RNASeq findings. All-stage Sertoli marker *SOX9* and BMP4 showed 46-fold and 3.5-fold increases in 6TF SLCs over GFPC, respectively (Figure 2-figure supplement 1F). Immature marker *PTGDS* showed a 2-fold increase on linear scale. 6TF SLC showed no change in germ cell marker *DDX4* or in granulosa marker *FOXL2* and myeloid marker *ACTA2*. (Similar results were found for 8TF; Figure 2-figure supplement 1F).

Finally, the molecular phenotype was ascertained at the protein level. Sertoli cells express the transcription factors SOX9 and DMRT1 in the nucleus across all stages and in the immature stage respectively (Makoto Ono & Harley, 2013). Sertoli cells also exhibit cytoplasmic expression of signaling molecules AMH during the immature stage and DHH in all stages (Table S1). Immunofluorescent imaging revealed that all these four markers exhibited Sertoli-specific sub-cellular expression in 6TF SLCs (Figure 2E) as well as in 8TF SLC (Figure 2-figure supplement 2 A,B,C,D). The percentage of the cell population showing these appropriate expression patterns was higher in 6TF than in 8TF (Figure2-figure supplement 2G) confirming that 6TF was the better transduction condition.

### Trans-differentiation strategy applied to dermal fibroblasts of different 46, XY DSD genetic backgrounds and 46,XX produced SLCs that exhibit morphological changes

The method described here was able to trans-differentiate dermal fibroblasts derived from a 46, XY adult male into Sertoli-like cells. To test the robustness of the method, it was applied to commercially available dermal fibroblasts from individuals with various DSD: 46, XY SRXY1 (46,XY female “sex reversal” of unknown genetic etiology; no variant in known DSD genes was found by exome sequencing, not shown), 46,XY CD (Campomelic Dysplasia; a large heterozygous deletion including *SOX9* was identified by whole genome sequencing, not shown); 46, XY WT1 (Wilms’ Tumor; a typical large deletion including the *WT1* gene was demonstrated by the company), as well as a 46,XX healthy female.

These four cell lines were separately seeded, transduced and cultured in the same way as described previously to produce 6TF and 8TF SLC and GFPC for each line. The live cultures were imaged on Day 7 and Day 28 of the culture under the GFP filter and the images were analysed for changes in shape (shape factor) and size (area).

The 6TF SLCs for three 46, XY DSDs showed significant changes in shape (Figure 3A,B,C) and size (Figure 3-figure supplement 1 A,B,C) over the 1-month differentiation period. In contrast, the 46, XX 6TF SLC did not exhibit a significant shape change but did show a size increase like others (Figure 3D, Figure 3-figure supplement 1D). [Data for 8TF are shown in Figure 3-figure supplement 1 A,B,C,D].

**Figure 3:**
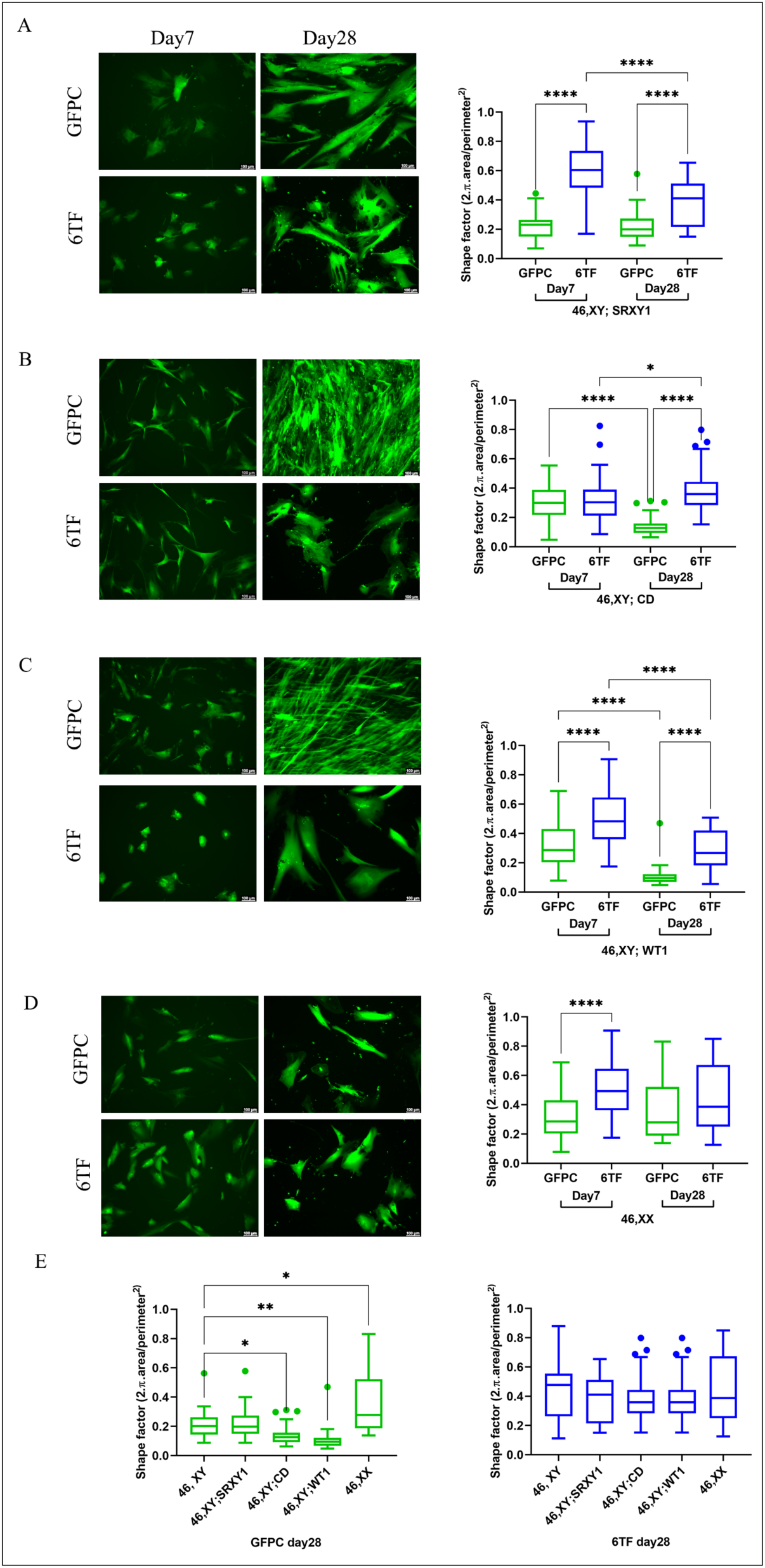
46,XYDSD and 46,XX fibroblast-derived SLCs exhibit varying degrees of change in morphology after 1 month trans-differentiation. Representative images of GFPC and 6TF SLCs on Day7 and Day28; Morphometric analysis of cells showing shape factor quantification represented as Tukey’s box plot with outlier values at either ends of the box (matching colored shapes), drawn from an average of 3 biological replicates (N), counting ~50-60 cells (n) within each group. * represent p values calculated from Mann-Whitney statistical tests conducted between the indicated groups. p > 0.05 (non significant, omitted from the graph), p ≤ 0.05 (*), p ≤ 0.01 (**), p ≤ 0.001 (***), p ≤ 0.0001 (****). (A) 46, XY; SRXYl (B) 46,XY; CD (Campomelic dysplasia) (C) 46,XY; due to WTl deletion and (D) 46,XX (E) Shape factor quantifications comparing GFPC (in green, left) and 6TF SLC (in blue, right)

We also compared these parameters in 1-month-old experiment groups of all these DSD and female lines with that of 46,XY. While the GFPC of the four lines had significantly different shapes at the start of the culture, these differences ebbed after the 6TF trans-differentiation process (Figure 3E). This interestingly was coupled with persistent differences in sizes of XY DSD and XX-derived GFPC and 6TF cells with those of 46,XY (Figure 3-supplementary figure 1E). This showed that cells derived from DSD individuals can also undergo transdifferentiation but with different characteristics than the control 46,XY cells and each other, possibly reflecting the different etiology of the DSD condition.

### Adhesion and proliferation of SLCs from XY, XX, and XY female DSD fibroblast lines

A key feature of Sertoli cells is their ability to adhere to form tubules then proliferate. SLCs derived from 46,XY, 46,XY with CD and 46,XX were subjected to xCELLigence Real Time Cell Analysis (RTCA) assays to measure their adhesion and proliferation properties. 0-6 hrs covers the time for adhesion of SLC suspensions to culture surface, while 6-48 hrs covers the period for cell proliferation. The Cell Index (CI) was plotted over time (Figure 4-supplementary figure 1 B,C,D,E,F) and the slope of CI curve was used as a measure to discern the differences or similarities between different experimental groups in their adhesion and proliferation phenotypes. The 46, XY 6TF SLCs showed lower adhesion (Figure 4A,) and slower proliferation (Figure 4B) than the GFP control. This might be indicative of it cell differentiation, which usually correlates negatively with proliferation. By contrast, 46 XY 8TF SLCs exhibited higher adhesion than GFPC while showing slower proliferation (Figure 4-supplementary figure 1A).

**Figure 4:**
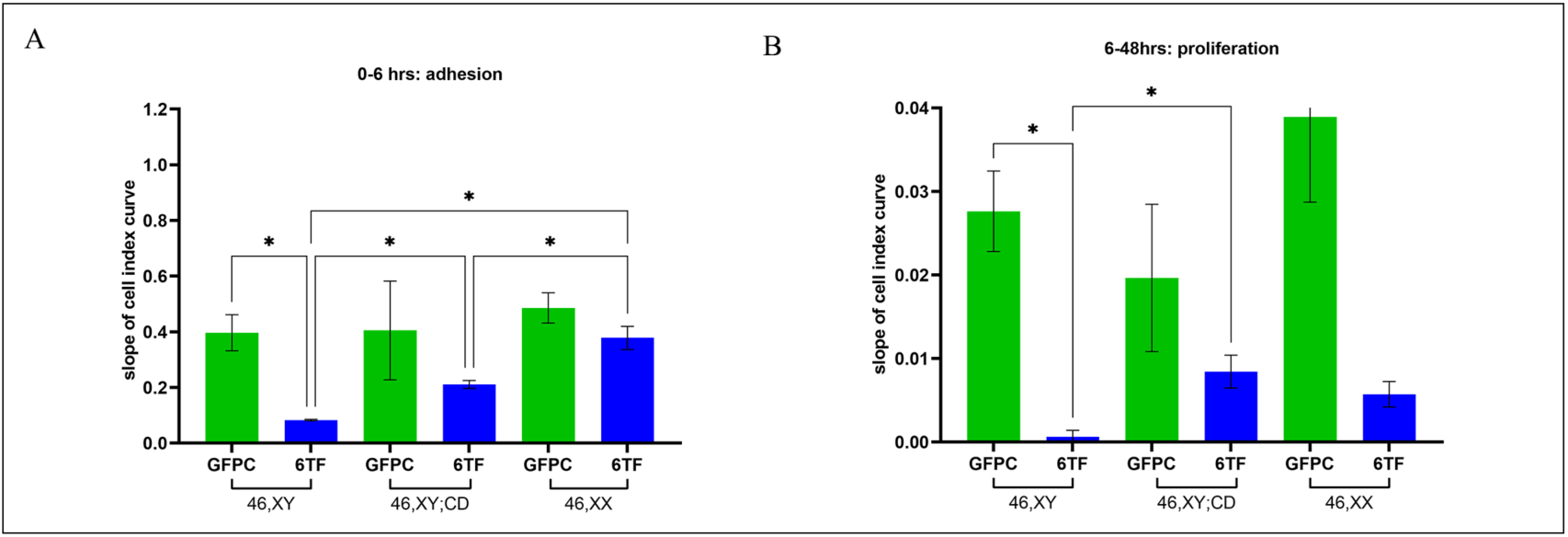
xCELLigence assay shows that 6TF SLCs exhibit distinct adhesion and proliferation phenotype than GFP controls across different genetic backgrounds. Slope of 0-6 hours cell index (CI) adhesion curves (A) and slope of 6-48 hrs cell index (CI) proliferation curves (B) for 1 month old GFPC and 6TF SLC derived from fibroblasts of indicated genetic backgrounds. all the experimental readings represent an average of three biological replicates (N=3); p > 0.05 (non significant,omitted from the graph), p ≤ 0.05 (*), p ≤ 0.01 (**), p ≤ 0.001 (***), p ≤ 0.0001 (****)

We also analyzed the adhesive and proliferative abilities of SLCs derived from the 46, XYwith CD and 46,XX cells. The GFP control fibroblasts of all the three cell lines showed similar adhesion and proliferation profiles (Figure 4A,B) despite their genetic differences. Sex dimorphism was observed in the adhesive abilities of the 46,XY SLCs and 46,XX SLCs, with the 46,XX SLCs showing a significantly higher adhesive ability (Figure 4A). The 46,XY;CD SLCs showed an intermediate adhesive ability, significantly higher than the 46,XY SLCs but significantly lower than the 46,XX SLCs. A trend towards sex dimorphism was also observed when comparing the proliferation of 46,XY SLCs and 46, XX SLCs although this did not reach statistical significance (Figure 4B). The 46,XY;CD SLCs once again showed significantly higher proliferation than the 46,XY SLCs, to a comparable level with the 46,XX SLCs (Figure 4B).

### 46,XX and 46,XY DSD fibroblasts derived SLC exhibited altered levels of Sertoli marker gene expression

The 46,XX- and 46,XY DSD-derived SLCs were evaluated for expression of Sertoli cell markers. Principal component analysis (PCA) of transcriptomes showed that all the samples belonging to each of the four cell lines clustered away from aSC as observed earlier for 46,XY SLCs (Figure 5-figure supplement 1). The 6TF and 8TF samples clustered away from GFPC in 46, XY; SRY1, 46,XY; WT1 and 46, XX but they didn’t separate as much in 46,XY;CD (Figure 5-figure supplement 1 A,B,C,D). The correlation plot for GFPC, 6TF SLC and 8TF SLC derived from each of the four cell lines showed that the replicate samples correlated with each other and each of the three experimental groups within each line were distinct from one another (Figure 5-figure supplement 2 B,C,D,E). Together these data suggest that the experimental procedure had a high level of reproducibility.

We investigated the fold change in expression of known DSD genes and Sertoli markers. Notably, while 46, XX SLCs expressed the expected Mogrify TFs in 6TF and 8TF SLCs (Figure 5-figure supplement 3D) they did not show increased *SOX9* expression over GFPC, either in the RNA seq data (Figure 5A) or in qPCR data (Figure 5-figure supplement 3 E). Even though they didn’t express *SOX9*, they did express bipotential gonad markers *GADD45G, ZFPM2, NR0B1* and immature SC markers *KRT18, INHBA, NCAM1/2, HSD17B3* and the all-stage marker *GDNF*. They barely expressed any mature markers, unlike 46,XY SLCs. Even though their SC marker expression profile was subdued in comparison to 46,XY SLCs, the presence of *NR5A1* and *EBF1* in the transduction mix had a similar effect on 8TF XX SLC in inducing Leydig markers *HSD3B2* and *CYP11A1* (Figure 5-figure supplement 3 D). Lack of induction of *SOX9* in 46,XX derived SLCs interestingly coincides with lack of significant morphometric change in their SLCs (Figure 3D).

**Figure 5:**
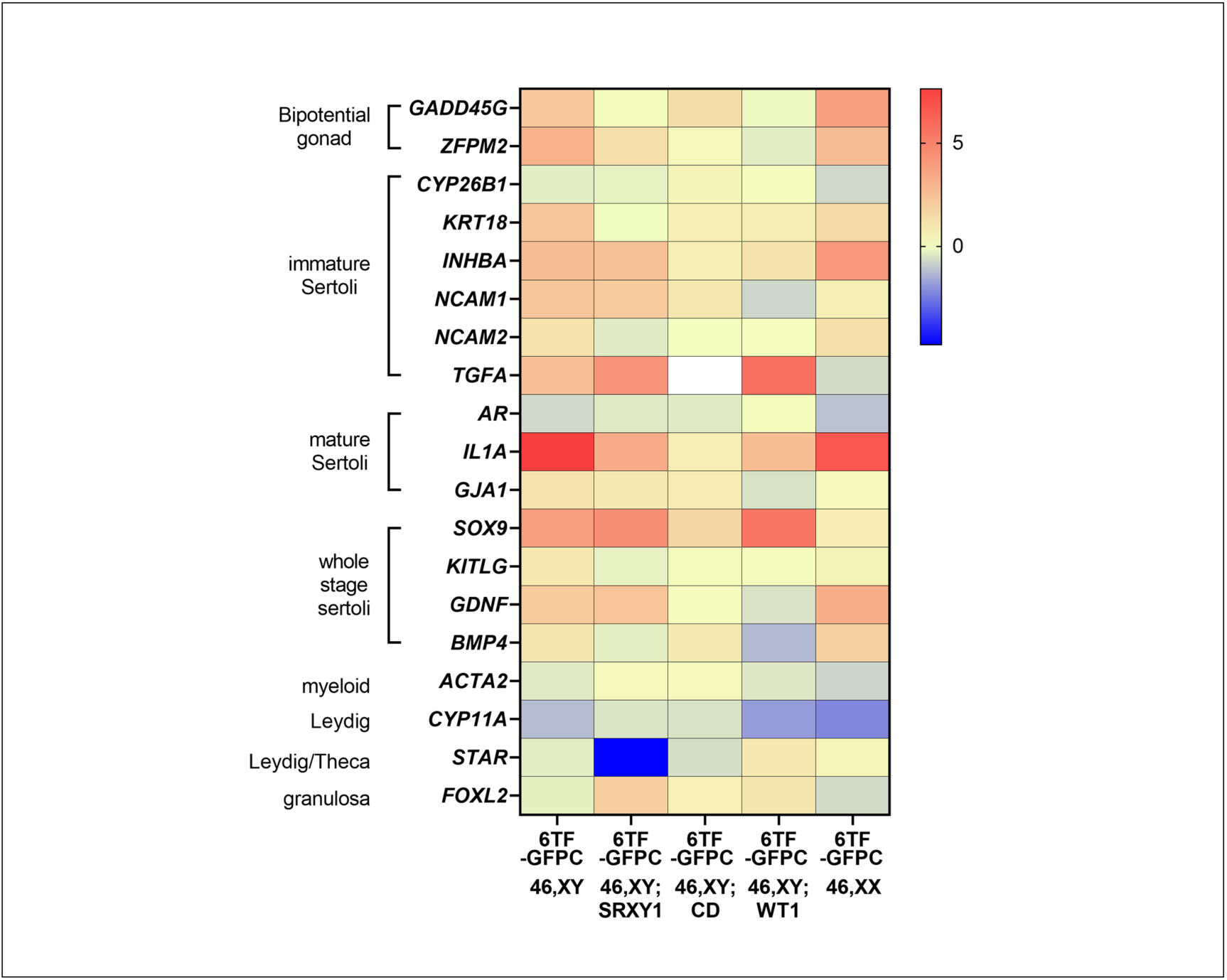
6TF SLCs derived from 46,XY DSD and 46,XX fibroblasts showed varying levels of Sertoli cell marker gene expression. Heat map showing 6TF vs GFPC log2 fold change for indicated gonadal marker genes as measured by RNAseq for each of the indicated genetic backgrounds: 46,XY (N=3), 46,XY; SRXY1(N=2); 46,XY; CD (N=3), 46,XY; WTl (N=3), 46,XX (N=3).

6TF and 8TF SLCs derived from 46, XY, SRXY1 DSD cells showed induction of *SOX9* by both RNAseq (Figure 5A) and qPCR (Figure 5-figure supplement 3E) as observed in 46,XY-derived SLCs. These SLCs exhibited similar profiles of expression of bipotential gonad markers *LHX9, GADD45G, ZFPM2*; immature SC markers *KRT18, INHBA/B, NCAM1* and all-stage marker *GDNF*. This showed that the trans-differentiation method was capable of pushing cells from DSD background genotype towards a Sertoli fate.

46, XY, CD derived 6TF SLCs showed increased *SOX9* expression (Figure 5A; 8TF shown in 5-figure supplement 3 B,E) and a modest, not statistically significant rise in the expression of *PTGDS, NCAM1/2, INHB* and *BMP4*. *SOX9* was also expressed in 46, XY, WT1-derived 6TF and 8TF SLCs at levels comparable to 46, XY-derived SLCs (Figure 5A, 5-figure supplement 3 C,E). These SLCs expectedly exhibited an increase in expression of other immature SC markers like *CYP26B1, PTGDS, KRT18, INHBA* and *NCAM2*. They additionally expressed mature markers *CLDN11, CTSL* and another all-stage marker *BMP4.* 8TF SLCs derived from all the three DSD fibroblasts also exhibited higher levels of expression of Leydig markers *HSD3B2* and *CYP11A1* in comparison to 6TF SLCs, just like observed for 46,XY SLCs.

The transcriptomic profiling of these cells shows that the trans-differentiation method is robust enough to bring about fibroblast to Sertoli cell fate differentiation in cells belonging to different genetic backgrounds with some properties sexually dimorphic and DSD SLCs showing intermediate, distinct phenotypes.

## Discussion

Cell differentiation can be achieved by utilizing a transgene-free approach of supplementing the culture media with suitable growth and differentiation supplements. Sertoli and gonadal cell types have been produced using such approaches from different cell types like human embryonic stem cells (hESC), human induced pluripotent stem cells (iPSC) and human umbilical cord-derived perivascular cells (HUCPVC) (Bucay et al., 2008; Knarston et al., 2020; Rodríguez Gutiérrez, Eid, & Biason-Lauber, 2018; Shlush et al., 2017). A major drawback of such an approach is that they are transient, and target differentiation fate may not pass down to the daughter cells optimally, thus limiting the scope of utilization of resultant cells. Creating a cell model to aid DSD diagnosis in a patient-specific manner warrants a transgene-based approach that to produce genetically stable Sertoli-like cells. One of the most critical considerations in such operations is getting the right combination of transcription factors (TFs) and culture conditions that could drive the targeted cell fate change. Murine and human fibroblasts have been trans-differentiated into Sertoli-like cells using such a transgene approach. Murine fibroblasts were reprogrammed to a Sertoli fate using transgenic expression of *Nr5a1*, *Wt1*, *Dmrt1*, *Gata4*, and *Sox9* (Buganim et al., 2012). Transgenic expression of *NR5A1* and *GATA4* in human fibroblasts achieved expression of Sertoli markers, but specificity wasn’t clearly reported (Liang et al., 2019), and collateral induction of other gonadal cell types could not be ruled out.

Computational algorithms using published literature and datasets to predict TFs required for directed cellular reprogramming have been successfully used for other cell types (Alessio et al., 2015; Morris et al., 2014; Rackham et al., 2016). We utilized one such predictive tool, Mogrify, which suggested 8 or 6 TFs as being sufficient to transdifferentiate dermal fibroblasts to Sertoli cells (Figure 1): *FGF2, GATA6, GATA4, MXI1, JUNB, NFYB + NR5A1* and *EBF1,* This method, when applied on 46,XY fibroblasts, produced Sertoli-like cells that exhibited significant change in morphology (Figure 1), as well as expression of Sertoli molecular signatures as measured by RNAseq, qPCR as well as IF (Figure 2, Figure 2 supplement), and cell behavior (Figure 4). They also exhibited upregulation of several genes associated with known Sertoli phenotype including cell adhesion, proliferation, and lipid metabolism as revealed by GO analysis of co-DEG between them and an adult Sertoli transcriptome (Figure 2-figure supplement 2). Both 6TF and 8TF SLCs exhibited predominantly immature Sertoli molecular signatures rather than mature SC signatures (Figure 2). We found that the 6TF trans-differentiation condition was best suited to create SLCs, as the additional *NR5A1* and *EBF1* in 8TF conditions induced expression of unwanted Leydig cell markers (Figure 2-figure supplement 1). The method, when applied to fibroblasts of different 46, XY DSD and 46, XX genotypes, produced varying levels of success in achieving the Sertoli phenotype. While 46, XY; SRXY1 and WT1 SLCs achieved comparable levels of change in morphometry and Sertoli marker expression, 46, XY; CD and 46,XX SLCs exhibited significantly less change than was observed in 46, XY SLCs (Figure 3, Figure 5, Figure 5-figure supplement 3). When comparing adhesion and proliferation of SLCs derived from different backgrounds also, the 46, XX SLCs showed quite a difference from 46, XY SLCs whereas 46,XY CD SLCs were somewhat in between (Figure 4). The varying levels of success in achieving that phenotype in different DSD cells might be reflective of their different genetic backgrounds and of which level of the genetic networks are affected by the mutations carried by those DSD cells. This trans-differentiation method can serve as a robust resource for precision medicine as it can be employed to transduce DSD patient-derived fibroblasts into Sertoli cells, which in turn can be used for relevant gonadal developmental stage specific transcriptomic and phenotypic analysis, thus aiding DSD molecular diagnostics.

## Materials and Methods

### Plasmids, lentiviral particle generation and titration

Second-generation lentiviral packaging plasmid pMD2.G encoding HIV-1-derived gag (structural protein), pol (retroviral enzyme), tat (transcriptional regulator) and rev (post-transcriptional regulator) and the envelope plasmid psPAX2 encoding VSV-G coat protein were purchased from Addgene repository. Eight different plasmids with human *FGF2, GATA6, GATA4, MXI1, JUNB, NFYB, NR5A1* and *EBF1* respectively, cloned into the pEZ-Lv165 lentiviral transfer vector system were purchased from GeneCopoeia, Inc. (Rockville, MD 20850, USA). pEZ-Lv165 vector system allows for robust parallel expression of the cloned gene and GFP marker protein separated by IRES2 sequence under the control of a strong EF1-alpha bi-cistronic promoter and it lacks mammalian antibiotic selection.

HEK-293T cells were seeded at 2.5×10^6^ in 25ml of DMEM (Genesee Cat. 25-500) completed with 10% heat-inactivated FBS (Thermo Cat. 16000044) in five 150 mm tissue culture dishes (Millipore Sigma Cat. CLS430599) and cultured for three days at 37^0^C in humidified tissue culture incubator with 5% CO2 supply to reach 80% confluency. 2 hours before transfections, the media was replaced with 22.5 mL of fresh complete media. Plasmid transfection was carried out by the calcium chloride method. 112.5 μg of transfer vector plasmid with gene of interest, 39.5 μg of packaging plasmid and 73 μg of envelope plasmid were mixed in a conical tube. 3.3 mL of 0.1X Tris-EDTA pH 8.0, 1.75 mL of 2.5 mM HEPES buffered distilled water pH 7.3, and 565 μL of 2.5M CaCl2 was added and mixed well by pipetting. 5.7 mL of 2x HEPES buffered saline (0.28 M NaCl, 50 mM HEPES, 1.5 mM Na2HPO4 dissolved in dH2O and pH adjusted to 7.0 with 10N NaOH) was added dropwise to the mixture under vigorous agitation and incubated at room temperature for 15 minutes. 2.25 mL of this transfection mix was added into each of the five dishes, they were swirled gently to mix and put back into the incubator overnight/12 hours. Early the next morning, the transfection media in each dish was replaced with fresh 14 mL complete media for the transfected cells to generate lentiviral particles and release into the media. Three subsequent harvests of lentiviral particle-containing media were collected 8 hours, 12 hours and 8 hours apart from the five dishes and entire ~210 mL of the harvest was pooled together, stored at 4^0^C. The pooled harvest was centrifuged at 500 G for 5 min at 4^0^C to pellet the cell debris and filter sterilized using 0.22 microns filter cup (Sigma Millipore Cat. CLS430769). Equal volumes were distributed into 6 ultra-clear thin wall tubes (Beckman Coulter Cat. 344058) and ultra-centrifuged at 50,000 G for 2 hours at 16^0^C in a Beckman Coulter Optima XE ultracentrifuge to precipitate lentiviral particles. The media supernatant was aspirated out and the lentiviral pellet was left to dry at RT for 10 mins. The pellet in each of the six tubes was dissolved in 17.5 μL of DPBS (Genesee Cat. 25-508) by gentle pipetting, avoiding air bubble formation and pooled. The pooled concentrated lentivirus was divided into 20 µL aliquots and stored at −80^0^C. All the surfaces, used dishes and spent media were thoroughly decontaminated with bleach before discarding.

The lentivirus was titrated by estimating the GFP positive cell population using flow cytometry. 1×10^5^ HEK 293T cells were re-suspended in 0.5 mL of complete media and seeded into each well of a 12-well dish (Millipore Sigma Cat. CLS3513). On the next day, cells were transduced with serial dilutions of 1μL, 10^−1^μL, 10^−2^μL, 10^−3^μl and 10^−4^μL lentivirus mixed in 0.5 mL media and 0.5 μL of 100 mg/mL Polybrene (Millipore Sigma Cat. TR-1003-G) in duplicate wells. One of the wells was used to harvest and count cell numbers at the time of transduction (day 1 cell count). The dish was left to incubate for 3 days, the transduced cells were harvested by trypsinization and fixed in 1% formaldehyde-DPBS for 5 mins at RT. They were washed once with DPBS and finally re-suspended in DPBS. The samples were analyzed on a Beckman Coulter Cytoflex S flow cytometer instrument for green fluorescence to determine GFP-positive cell percentage. Lentiviral dilution samples yielding 1-20% GFP positives were chosen for titer calculations as any percentage higher than that insinuates multiple expressible viral integration events per cell in the population and less than 1% can be erroneous/false positive. Viral titer concentration was calculated in transducing units / μl as (day 1 cell count x % GFP-positive cells/100)/volume of lentivirus in μl.

### Cell lines and trans-differentiation culture

Normal human adult 46, XY primary dermal fibroblast (HDFa) (Cat. ATCC PCS-201-012) was purchased from American Type Culture Collection repository (ATCC). The following cell lines were purchased from Coriell cell repository: GM03651: Skin fibroblasts derived from a 25-year-old 46, XX healthy female; GM00083: Gonadal fibroblasts derived from a 1-year-old 46, XY female; SRXY1; GM06938: fibroblasts derived from a 3-week-old male with a *WT1* deletion; GM21894: skin fibroblasts derived from a 46,XY patient with campomelic dysplasia; and GM03368: gonadal fibroblasts derived from a 17-year-old 46, XY woman). HDFa was cultured in Medium 106 (Thermo Cat.M106-500), completed with low serum growth supplement (LSGS) (Thermo Cat. S00310), and 1X penicillin-streptomycin antibiotic (P/S) (Thermo Cat. 15140122). GM03651, GM00083, GM06938 and GM12896 were cultured in Eagle’s minimum essential medium (Thermo Cat. M0325) completed with 15% non-inactivated FBS (Thermo Cat. 16000044) and 1X P/S (Thermo Cat. 15140122). GM21894 was cultured in DMEM (Genesee Cat. 25-500) completed with 10% non-inactivated FBS and 1X P/S. All these cells were grown as adherent monolayer cultures in tissue culture-treated T25 flasks (Genesee Cat. 25-207), harvested by trypsinization upon 80-90% confluence, quenched by trypsin neutralization solution (Thermo Cat. R002100), spun down at 150g for 4 min at RT and sub-cultured in 5ml of suitable complete media. The media was replaced every alternate day in all these cultures until they reached confluency.

For the trans-differentiation culture, fibroblasts were seeded at 2.25×10^4^ into each well of 6 well dishes (Millipore Sigma Cat. CLS3516) in 2ml complete M106 on Day 0 and grown overnight. On Day 1, cells were counted from an extra well to determine the number of lentiviral particles required to transduce the cells at a multiplicity of infection (MOI) of five. Three transduction sets were designed: A. GFP; B. 8 TFs: *FGF2*, *GATA6*, *GATA4*, *MXI1*, *JUNB*, *NFYB*, *NR5A1* and *EBF1*; C. 6TFs: *FGF2*, *GATA6*, *GATA4*, *MXI1*, *JUNB*, and *NFYB*. The old media in wells was replaced with transduction media consisting of 2 mL complete M106 mixed with an appropriate amount of respective lentivirus cocktail and 2 μL of 100 mg/mL Polybrene (Millipore Sigma Cat. TR-1003-G). The cells were left to transduce for 12 hours/overnight. On Day 2, transduction media was replaced with fresh complete M106 and cultured in it for 4 days with alternate days media change. On day 5, M106 was replaced with 2 mL of trans-differentiation media consisting of basal Sertoli Cell Media (SerCM) completed with 5% FBS, 1X Sertoli cell growth supplement (SerCGS) and 1X P/S (ScienCell Cat. 4521). The culture was maintained in this media for another 24 days until day 30 with alternate days media change.

16 transduction conditions were designed for the pilot study. Conditions 1,2,3,4,9,10 exhibited low transduction efficiency while conditions 15, 16 showed high levels of post-transduction mortality (data not shown, Figure 1-figure supplement 1C). Conditions 5, 6, 7, 8, 11, 12, 13, and 14 showed fairly high levels of transduction efficiency, minimal cell mortality, and variable degrees of morphological change in comparison to their respective GFP controls (Figure 1-figure supplement 1C). Appearance of a few Sertoli markers were explored using qPCR. Mature stage Sertoli marker *AR* (Table S1) didn’t appear in any of the conditions while all-stage marker *BMP4* appeared in conditions 5-8 and 13-16 (Figure 1-figure supplement 1B). Immature stage Sertoli marker *PTGDS* appeared in conditions 5, 6, 7, 8 while critical all-stage Sertoli marker *SOX9* appeared to be significantly expressed in conditions 5, 6 and to a lesser extent in 7, 8. (Figure 1-figure supplement 1B).

Stable viral transductions were confirmed by upregulation of each of the 8 TFs or 6 TFs after 1-month trans-differentiation culture. The 8TF and 6TF SLCs exhibited well over 1.5 fold increase for each of the 8 or 6 TFs. Furthermore, *NR5A1* and *EBF1* didn’t show an increase in 6TF SLCs, showing specificity of transduction (Figure 2-figure supplement 1E; Figure 5-figure supplement 3 A,B,C,D).

### RT-PCR, qPCR, and target gene expression fold change analysis

Cells were lysed and total RNA was isolated using RNeasy micro kit (Qiagen Cat. 74004) following the manufacturer’s instructions. All the qPCR and RT-PCR reagents were purchased from Thermo Scientific. Total RNA was reverse transcribed into cDNA by RT-PCR using High-Capacity cDNA Reverse Transcription Kit (Cat. 4368814) following the manufacturer’s instructions on a BioRad C1000 thermal cycler. The following thermocycling conditions were used: 25^0^C-10 min, 37^0^C-120 min, 85^0^C-5min, 4^0^C-hold. The cDNA was used to perform quantitative PCR using Taqman chemistry reagents. qPCR reactions were set up in triplicates using 5 ng of cDNA per reaction using TaqMan™ Fast Advanced Master Mix (Cat. 4444556) with TaqMan™ Gene Expression Assay-VIC probe (Cat. 4448490 Assay ID: Hs02758991_g1) for *GAPDH* housekeeping gene and respective target genes. Following TaqMan™Gene Expression Assay (FAM) target gene probes were used: *SOX9* (Cat. 4331182 Assay ID: Hs00165814_m1), *BMP4* (Cat. 4331182 Assay ID: Hs00370078_m1), *PTGDS* (Cat. 4331182 Assay ID: Hs00168748_m1), *AR* (Cat. 4331182 Assay ID: Hs00171172_m1), *HSD3B2* (Cat. 4331182 Assay ID: Hs00605123_m1), *DDX4* (Cat. 4331182 Assay ID: Hs00987125_m1), *FOXL2* (Cat. 4331182 Assay ID: Hs00846401_s1), *ACTA2* (Cat. 4331182 Assay ID: Hs00426835_g1). 12 μL reactions were set up following manufacturer’s instructions in MicroAmp™ EnduraPlate™ Optical 384-Well Clear GPLE Reaction Plates with Barcode (Cat. 4483319) on a cooling block, sealed, spun down briefly and subject to PCR on a QuantStudio™ 7 Flex Real-Time PCR platform (Cat. 4485701) using manufacturer suggested default thermocycling conditions for 40 amplification cycles. The target gene expression fold change in experimental over control was calculated using inverse comparative threshold cycle (ΔΔCt) method after normalizing for *GAPDH* as the internal loading control.

### Live cell imaging and morphometry

GFP, 8TF and 6TF transduced 1-month cultures growing in 6-well dishes and expressing GFP were imaged under the green fluorescence channel of a Leica DMIL LED Inverted Fluorescence Microscope using Leica’s LasX imaging software. The images were exported in TIFF format and used for morphometric analysis. Morphometry was performed using MetaMorph software. The freehand tool in the software was used to draw the outlines of GFP expressing cells, which then computed the perimeter as well as area enclosed within that cell perimeter in arbitrary units.. An average of 60-80 cells were counted per group and the group statistics were compared using unpaired t test. Cell surface area measured in arbitrary units was used as a readout for cell size as these are two-dimensional adherent cell cultures, while an arbitrary index called shape factor, defined as 2.π.area/perimeter2 was utilized as a proxy measurement of cell shape

### Immunofluorescence staining and imaging

For immunofluorescence staining, the cells were fixed with 4% paraformaldehyde for seven minutes and washed three times with phosphate-buffered saline (PBS). In brief, slides were blocked for 60 minutes with 1% donkey serum, followed by incubation with primary antibodies overnight at 4°C, including anti-SOX9 rabbit polyclonal (Sigma-Aldrich AB5535, 1:400), anti-GATA4 goat polyclonal (Santa Cruz SC-1237, 1:500), anti-DMRT1 rabbit polyclonal (Santa Cruz SC-98341, 1:200), anti-AMH mouse monoclonal (Santa Cruz SC-166752, 1:400), or anti-DHH goat polyclonal (Santa Cruz SC-1193, 1:300). After washing three times in PBS for five minutes each, the slides were incubated with the corresponding secondary antibody at a 1:1000 dilution (ThermoFisher A-21207, ThermoFisher A-11058, ThermoFisher A10037 or ThermoFisher A-31573.). DAPI (4,6-diamidino-2-phenylindole, Invitrogen) was used to label the nuclei. Cells were observed for epifluorescence under a Nikon C1 confocal microscope or Olympus FV1200 confocal microscope under standard conditions.

### RNA sequencing: Library preparation with polyA selection, HiSeq sequencing and data analysis

RNA library preparations and sequencing reactions were conducted at GENEWIZ, LLC. (South Plainfield, NJ, USA). RNA samples received were quantified using Qubit 2.0 Fluorometer (Life Technologies, Carlsbad, CA, USA) and RNA integrity was checked using Agilent TapeStation 4200 (Agilent Technologies, Palo Alto, CA, USA).

RNA sequencing libraries were prepared using the NEBNext Ultra RNA Library Prep Kit from Illumina using the manufacturer’s instructions (NEB, Ipswich, MA, USA). Briefly, mRNAs were initially enriched with Oligod(T) beads. Enriched mRNAs were fragmented for 15 minutes at 94 °C. First-strand and second-strand cDNA were subsequently synthesized. cDNA fragments were end-repaired and adenylated at 3’ends, and universal adapters were ligated to cDNA fragments, followed by index addition and library enrichment by PCR with limited cycles. The sequencing library was validated on the Agilent TapeStation (Agilent Technologies, Palo Alto, CA, USA), and quantified by using Qubit 2.0 Fluorometer (Invitrogen, Carlsbad, CA) as well as by quantitative PCR (KAPA Biosystems, Wilmington, MA, USA).

The sequencing libraries were clustered on a single lane of a flowcell. After clustering, the flowcell was loaded on the Illumina HiSeq instrument (4000 or equivalent) according to manufacturer’s instructions. The samples were sequenced using a 2×150bp Paired-End (PE) configuration. Image analysis and base calling were conducted by the HiSeq Control Software (HCS). Raw sequence data (.bcl files) generated from Illumina HiSeq was converted into fastq files and de-multiplexed using Illumina’s bcl2fastq 2.17 software. One mismatch was allowed for index sequence identification.

### RNAseq Data analysis

BCL files generated from sequencer were converted using bcl2fastq (Illumina) to fastq files, quality-checked using *fastqc* (https://www.bioinformatics.babraham.ac.uk/projects/fastqc/). Adapter and quality trimming was performed using *trimgalore* (https://www.bioinformatics.babraham.ac.uk/projects/trim_galore/). Summary QC files for before and after quality trimming was performed using MultiQC (Ewels et al., 2016). Alignment to hg38 reference was performed using *STAR* (Dobin et al., 2013), followed by count estimation using *RSEM* (Li & Dewey, 2011). For differential expression analysis, read counts were estimated using *tximport* (Soneson et al., 2016), followed by normalization and differential expression analysis performed using *Deseq2* (Love et al., 2014). All significantly expressed genes were above a threshold of absolute log2 fold change of 1.5 and p-value < 0.05. Gene ontology was performed using *gProfiler* (Raudvere et al., 2019), where we extracted only biological processes. Visualization was performed using pheatmap and ggplot2 R packages.

### Adhesion and proliferation xCELLigence assays on SLCs

For the xCELLigence assays Sertoli-like cells were lifted using 0.05% trypsin and counted on a haemocytometer. An xCELLigence E-Plate (ACEA BioSciences, VIEW 96 PET plate) was prepared with 50 μL of complete growth media in each well. All conditions were run in technical triplicate. GFP controls, 8TF SLC and 6TF SLC derived from three cell lines - HDFa (46, XY; healthy male), GM21894 (46, XY; CD) and GM03651 (46, XX; healthy female) - were analyzed using this technique. The E-plate was engaged into the xCELLigence Real Time Cell Analysis (RTCA) Instrument (Agilent) and background measurement of the wells plus growth media was recorded before adding cells in 150 μL of media for each SLC condition (5 x 10^3^ cells/well). The

E-plates were re-engaged onto the xCELLigence analyzer and incubated for 48 h at 37°C in a 5% CO2 atmosphere. Impedance values were recorded every 2 minutes for the first 6 hours and adhesion data derived. Between 6 hours and 48 hours impedance measurements were recorded every 15 minutes to derive the proliferation date. The value of the slope, i.e. the change in adhesion/proliferation over time, was used for data analysis. Statistical analysis was performed in Graphpad Prism 5 using a Student’s t-test.

## Acknowledgements

None

## Supplementary Figures

**Figure 1-figure supplement 1:**
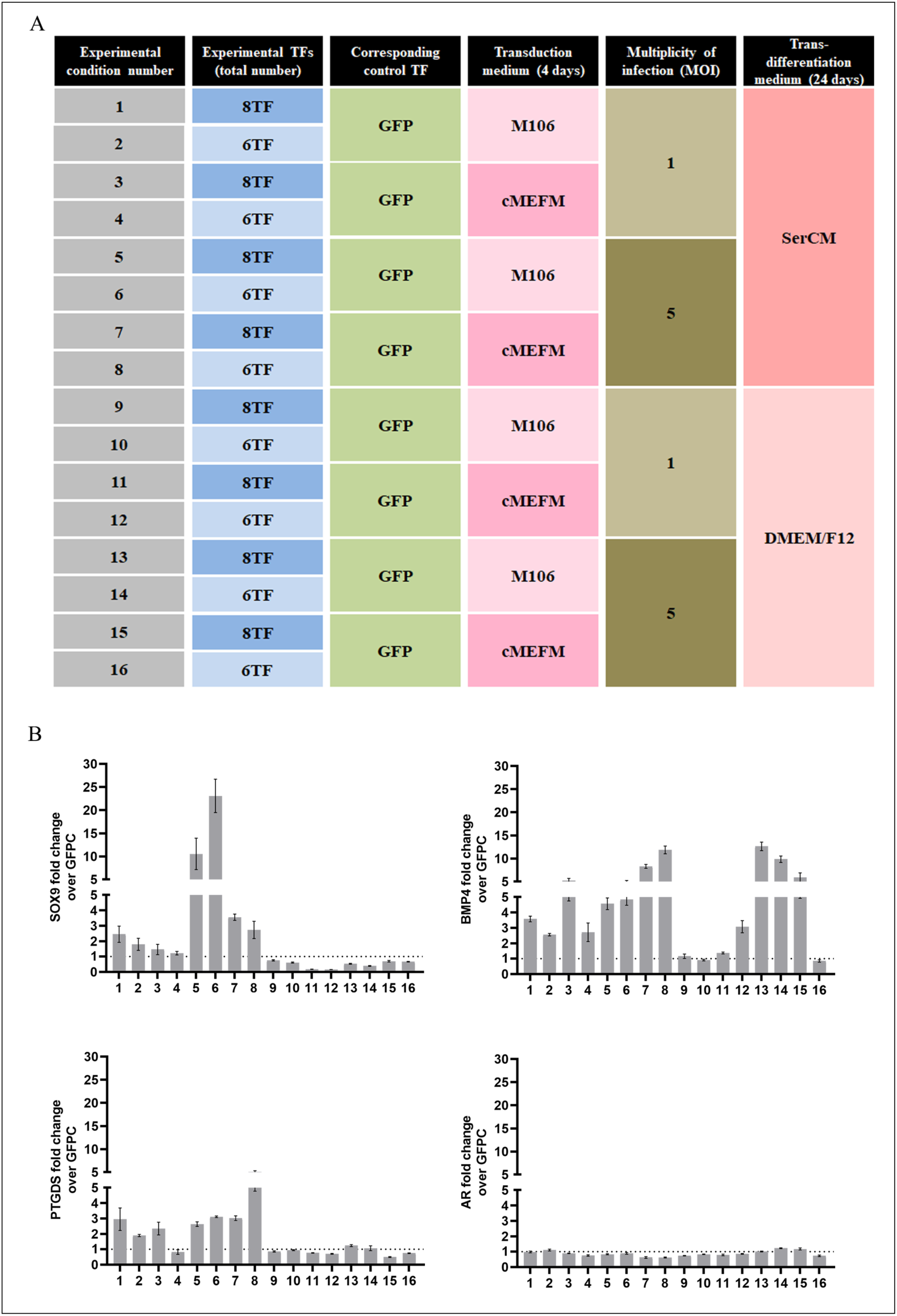
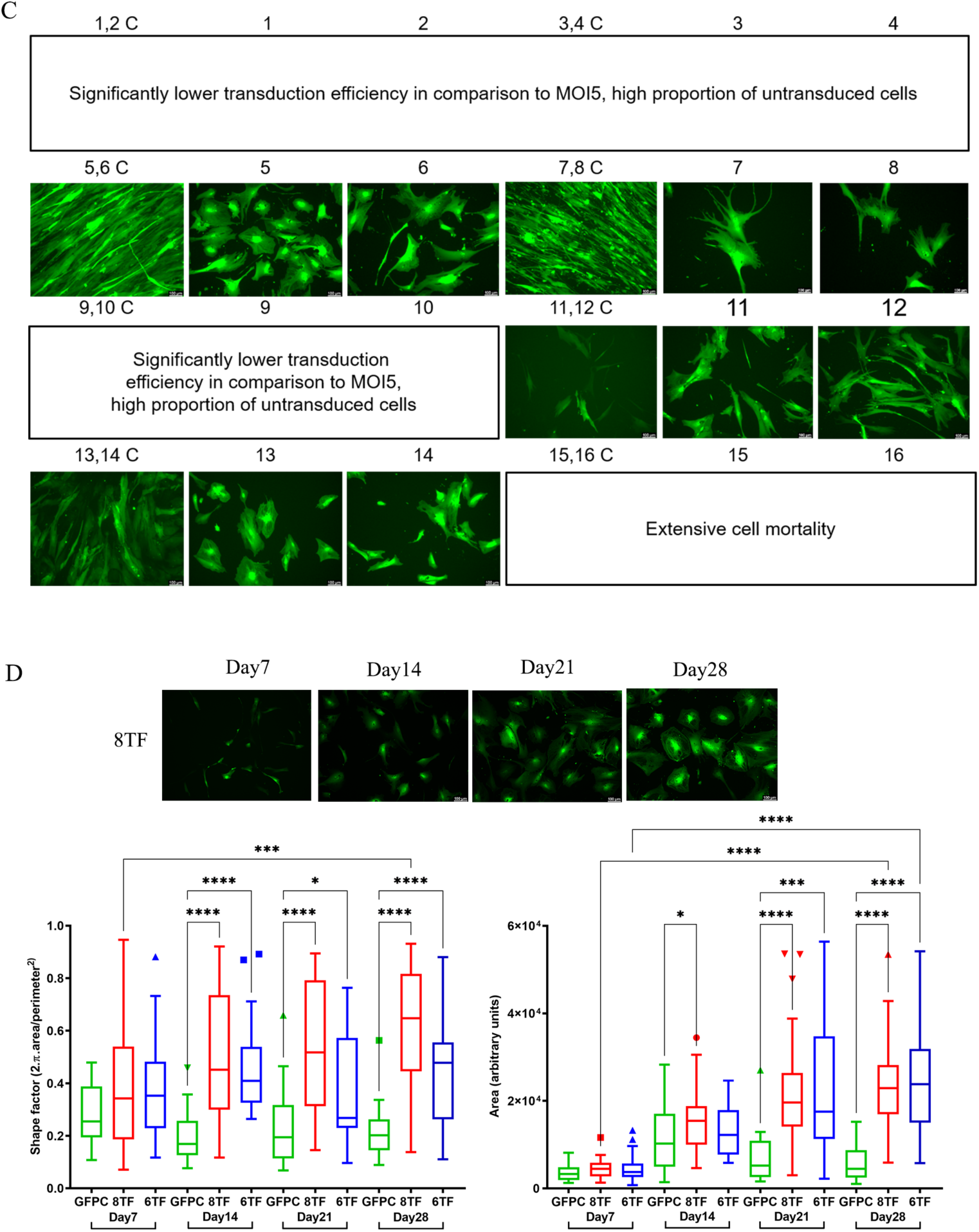
Standardization of trans-differentiation protocol. (A) Outline of the initial16 trans-differentiation conditions designed to attempt fibroblast-Sertoli cell like trans differentiations. (B)The outcome ofthese conditions was evaluated by looking for Sertoli marker expression in each ofthe 16 conditions by qPCR for: SOX9, BMP4, PTGDS andAR. N=2, n=6 for each of the qPCRs, error bars represent SEM. (C) representative images of 1 month live cultures for all these conditions. Missing images either had too less green fluorescence or had extensive cell death. (D) Representative images of 46,XY SLC culture over 1 month, Morphometric analysis of GFPC, 8TF and 6TF cells showing shape factor quantification represented as Tukey’s box plot with outlier values at either ends of the box (matching colored shapes), drawn from an average of 3 biological replicates (N), counting ~50-60 cells (n) within each group.* represent p values calculated from Mann Whitney statistical tests conducted between the indicated groups. p > 0.05 (non significant, omitted from the graph), p ≤ 0.05 (*), p ≤ 0.01 (**), p ≤ 0.001 (***), p ≤ 0.0001 (****)

**Figure 2-figure supplement 1:**
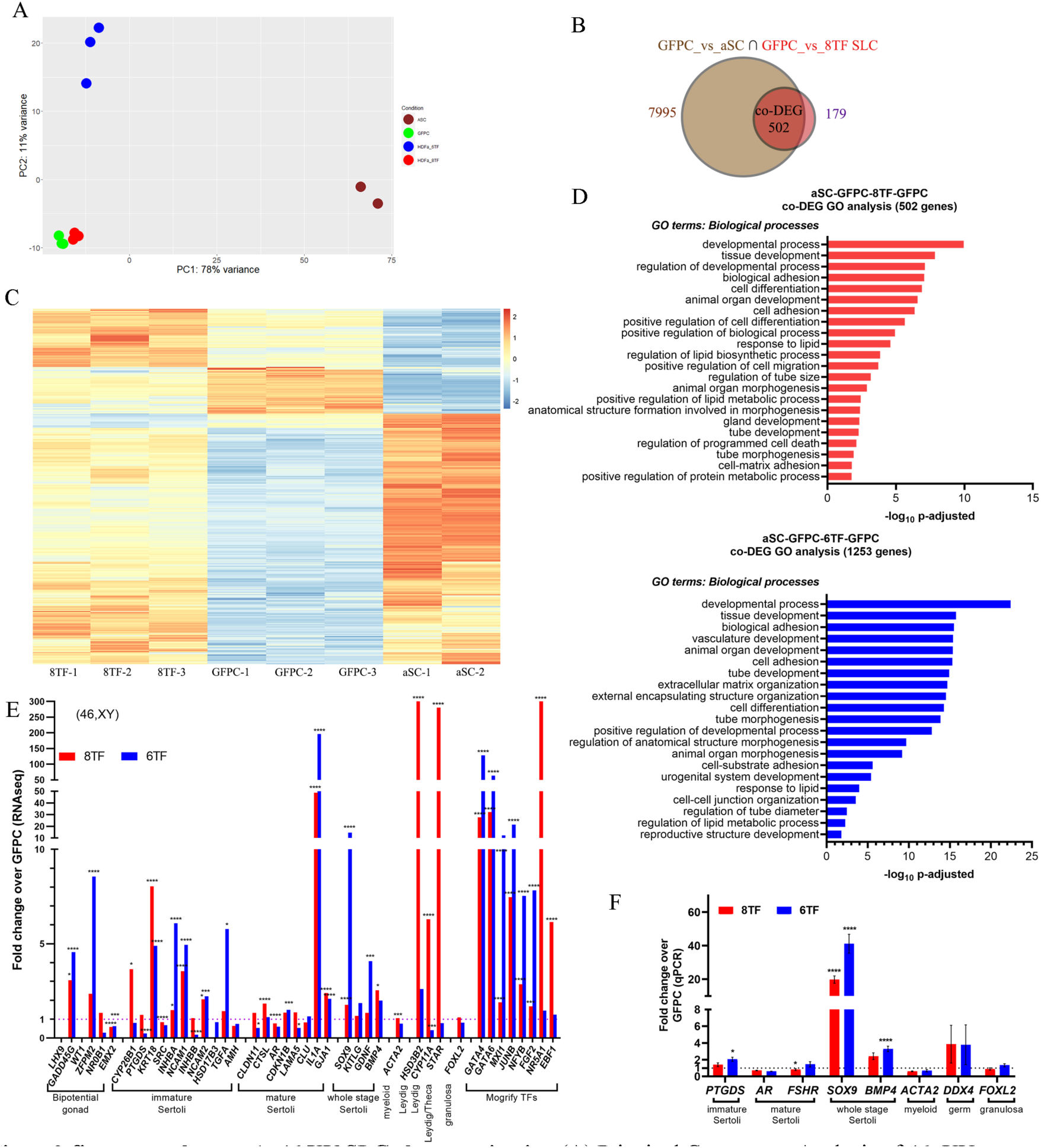
46,XY SLC cha’racterization. (A) Principal Component Analysis of 46, XY GFPC, 8TF, 6TF and adult Sertoli cells (aSC) transcriptomes. (B) Venn diagram displaying differentially expressed genes (DEGs, fold change>l.5, p adjusted <0.05) between GFPC (N=3) vs aSC (N=2) and GFPC (N=3) vs 8TF SLCs (N=3). Intersection part shows co-differentially expressed genes (co-DEGs). (C) Heatmap showing expression of co-DEGs for 8TF, GFPC and aSC (D) Biological process gene ontology (GO) analysis for the co-DEGs (E) RNAseq analysis showing linear scale fold change in expression of indicated markers in 46,XY derived 8TF and 6TF SLCs over GFPC N=3, p > 0.05 (non significant, omitted from the graph), p ≤ 0.05 (*), p ≤ 0.01 (**), p ≤ 0.001 (***), p ≤ 0.0001 (****)(F) Linear scale old change for indicated markers as determined by qPCR, the values represent mean+/- SEM of the following number of samples:PTGDS: N=9, n=18; AR: N=9, n=12; FSHR: N=4, n=16; SOX9: N=12,n=25; BMP4: N=9,n=18;ACTA2: N=3, n=3; DDX4: N=3, n=3; FOXL2: N=3, n=6. K: PTGDS: N=6, n=12; AR: N=6, n= 6; FSHR: N=9, n=15; SOX9: N=6, n=12; BMP4: N=6, n=12; ACTA2: N=3, n=3; DDX4: N=3, n=3; FOXL2: N=3, n=3. *represent p values calculated from unpaired t test conducted between each 8TF or 6TF and GFPC N=biological replicate, n=technical replicate

**Figure 2-figure supplement 2:**
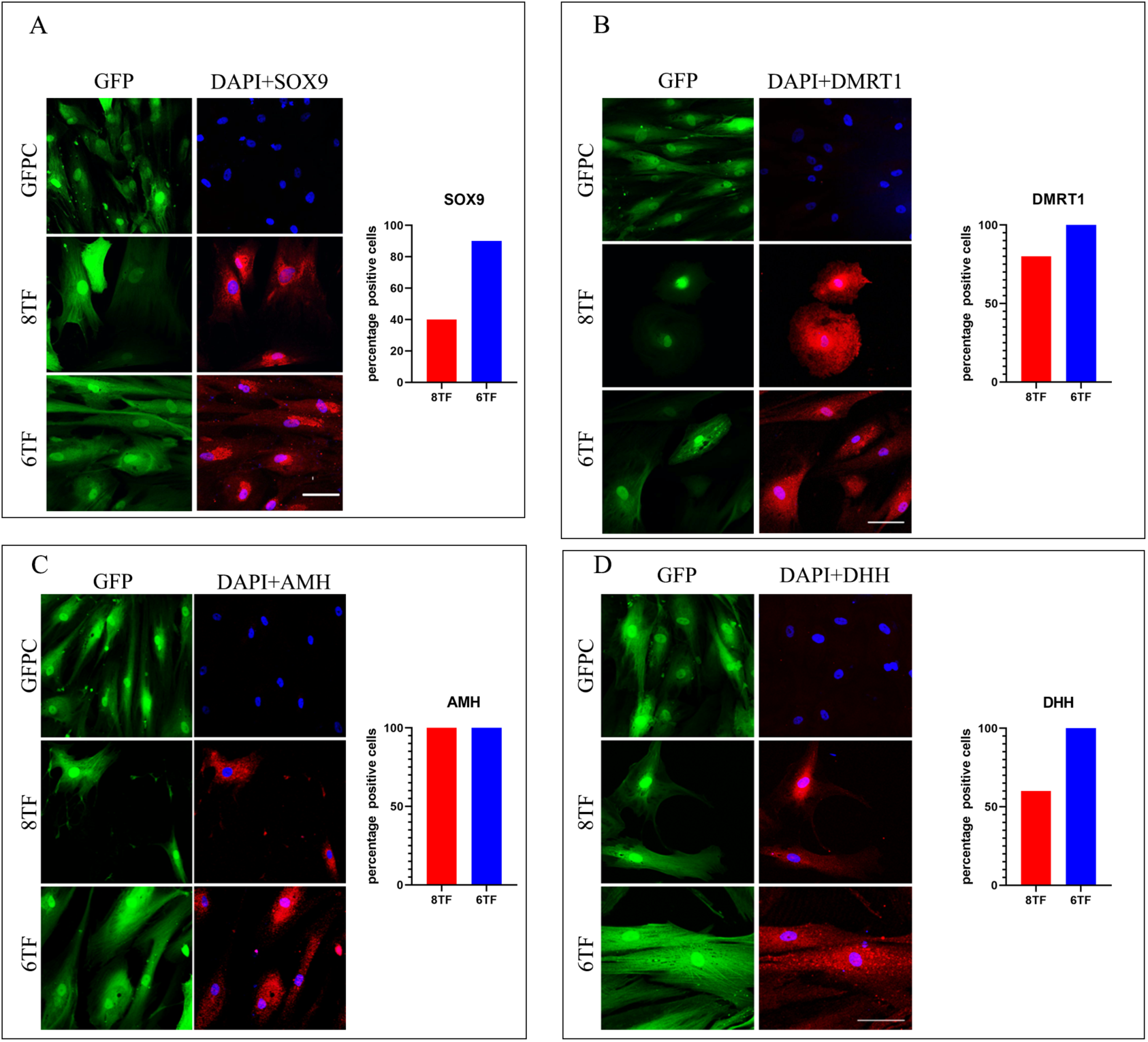
STF and 6TF SLCs show Sertoli specific marker expression. Representative immuno-fluorescence images showing subcellular expression and quantification of percentage of cells showing such expression (N=3, n=S0-60 for each group) for (A) SOX9 (B) DMRTI (C) AMH and (D) DHH in the 46, XY 8TF, 6TF SLCs and GFPC.

**Figure 3-figure supplement 1:**
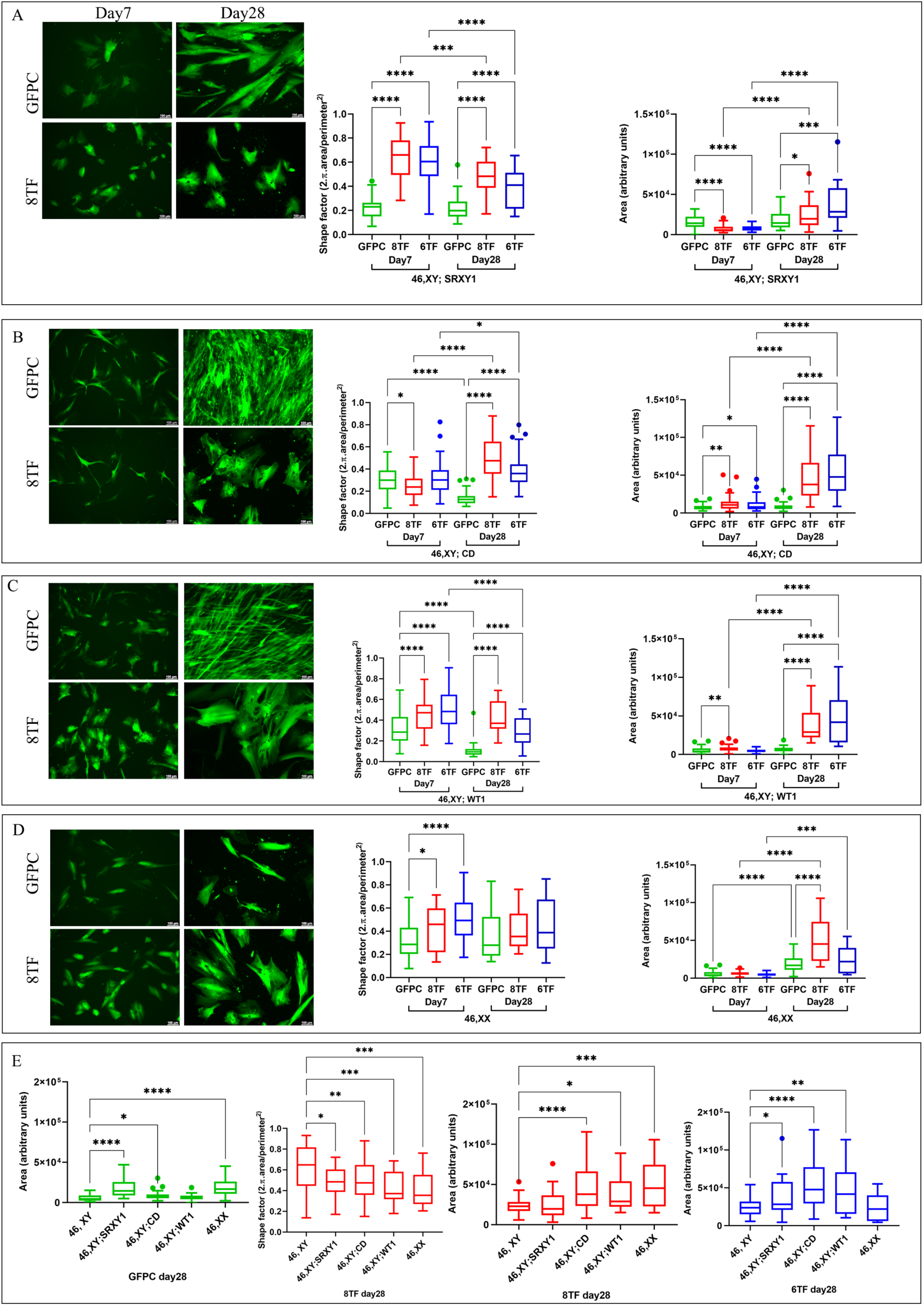
Morphometry of 46,XY DSD and 46,XX SLC: Representative images of GFPC and 8TF SLCs on Day7 and Day28; Shape factor and area quantifications for GFPC, 8TF and 6TF SLCs (N=3, n=50-60 for each group). * represent p values calculated from Mann-Whitney statistical tests conducted between the indicated groups for (A). 46, XY; SRXY1 (B) 46,XY; CD; (C) 46,XY; WT1 and (D) 46,XX; non significant p value comparisons are omitted (E) GFPC area factor, 8TF shape factor, 8TF area factor, 6TF area factor quantifications comparing 46, XY DSD and 46,XX genetic backgrounds with 46, XY.

**Figure 4-figure supplement 1:**
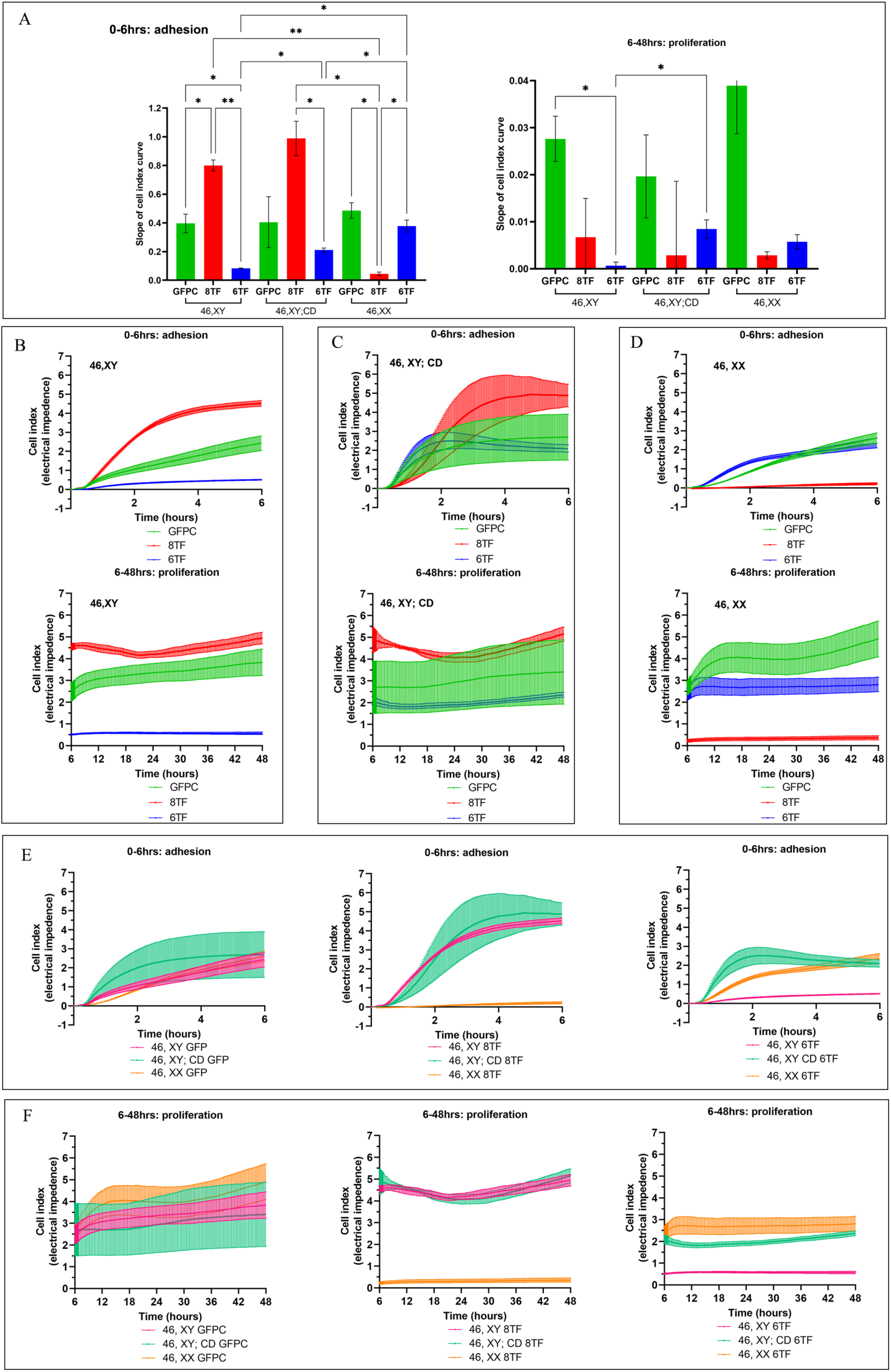
Cell index (CI) curves of xCELLigence assays. (A) Slope of 0-6 hours cell index (CI) adhesion and 6-48 hrs proliferation curves for 1-month-old GFPC, 8TF and 6TF SLC derived from fibroblasts of indicated genetic backgrounds. All the experimental readings represent an average of three biological replicates (N=3), * represent p values calculated from one way ANOVA test conducted amongst the three groups within each graph, ns is not shown. 0-6 hours CI adhesion curves and 6-48 hrs CI proliferation curves for 1-month-old GFPC, 8TF SLC and 6TF SLC derived from 46,XY (B), 46,XY;CD (C) and 46, XX (D). (E): Alternative representation of 0-6 hours cell index (CI) adhesion curves and (F) 6-48 hrs CI proliferation curves for 46,XY-, 46,XY with campomelic dysplasia (CD)-, or 46,XX-derived GFPC, 8TF SLCs and 6TF SLCs. All the data points represent an average plus SEM of three biological replicates (N=3).

**Figure 5-figure supplement 1:**
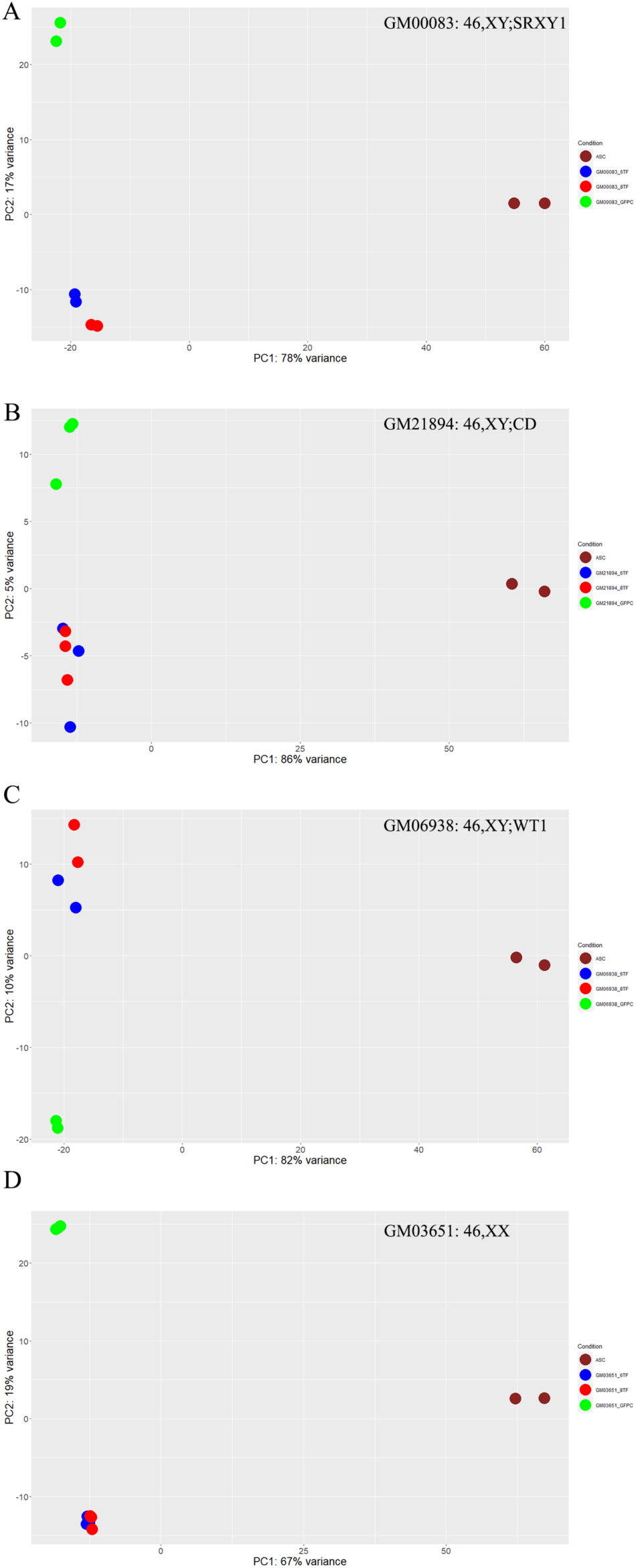
Principal Component Analysis of RNAseq samples. Principal component analysis(PCA) plots of RNAseq transcriptomes covering al1 the replicate samples for untransduced (abbrevated as ce11 line code_C), GFPC, 8TF, 6TF derived from (A) GM00083: 46,XY; SRXY 1, (B) GM 21894: 46,XY; CD, (C) GM0693 8: 46,XY; WT1 and (D) GM03 65 1: 46,XX along with aSC.

**Figure 5-figure supplement 2:**
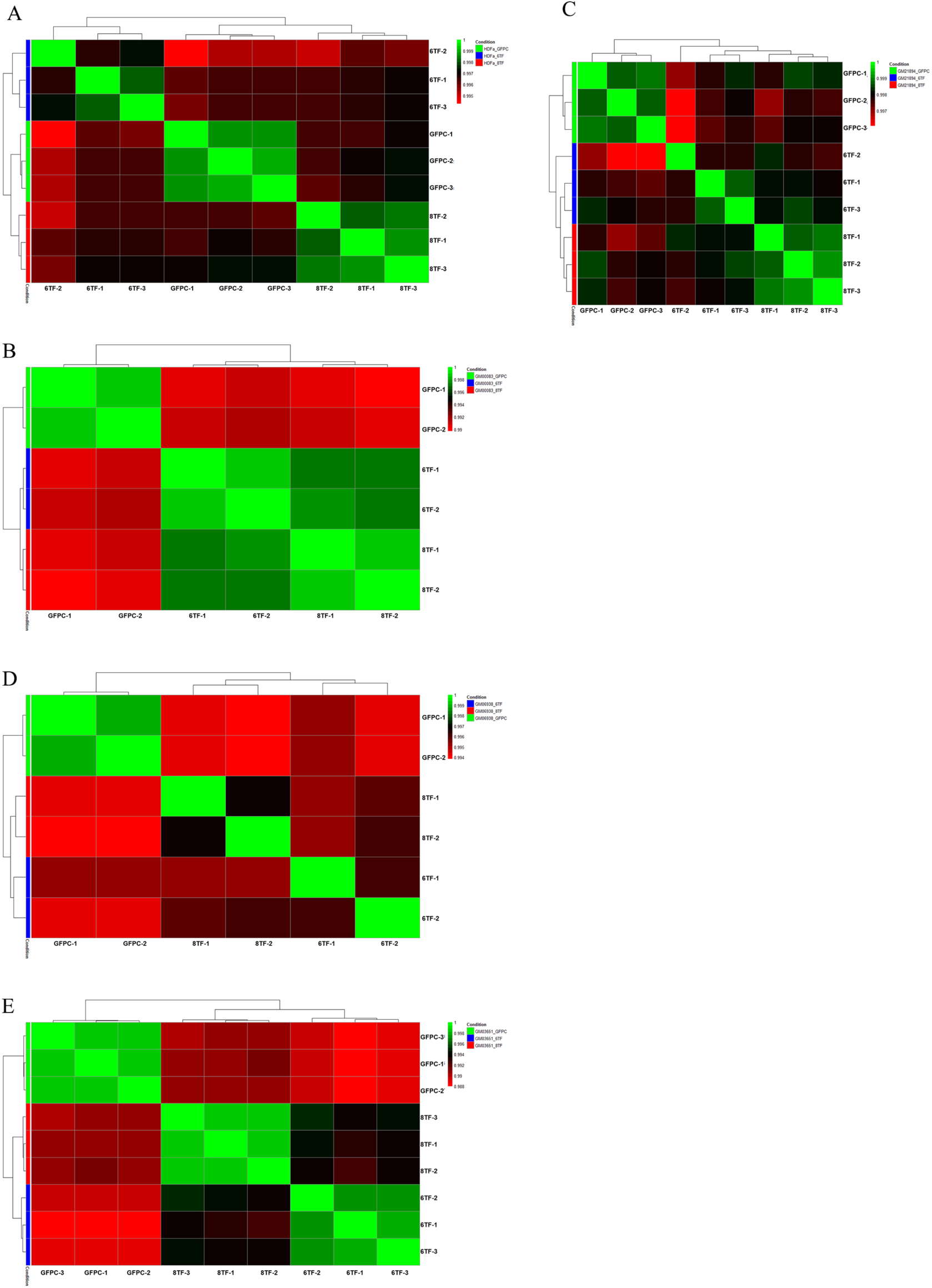
RNA seq sample replicates group together. Correlation plots for each of the 8TF, 6TF and GFPC RNA seq samples of (A) 46,XY (B) 46,XY; SRXYl (C) 46,XY; CD (D) 46,XY;WTl (E) 46,XX

**Figure 5-figure supplement 3:**
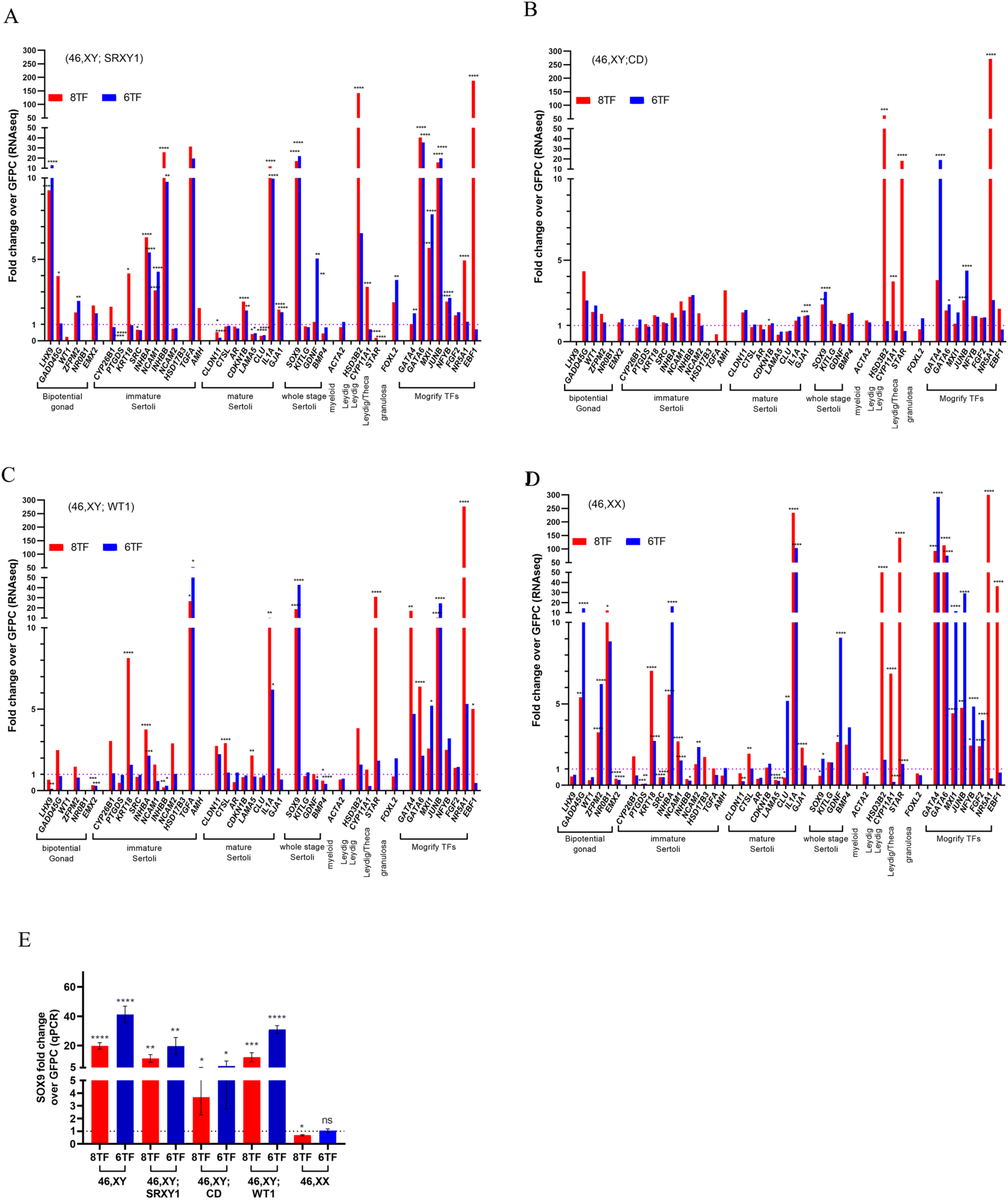
A wide range of gonadal markers and transduced Mogrify TFs show expression in SLC samples. RNAseq analysis shows linear scale fold change in expression of indicated markers in 8TF or 6TF derived SLCs in comparison to GFPC for (A) 46, XY; SRXYl (B) 46,XY;CD (C) 46,XY;WTl (D) 46,XX. N=3, p > 0.05 (non significant,omitted from the graph), p ≤ 0.05 (*), p ≤ 0.01 (**), p ≤ 0.001 (***), p ≤ 0.0001 (****). p value of fold change calculations between experimental and respective (E): Fold change for SOX9 as determined by qPCR, in both 8TF or 6TF SLC over respective GFPC for the indicated cell lines. N=5, n=25 for 46,XY, N=3,n=3 for 46,XY;SRXYl, 46,XY;CD, 46,CD; WTl and 46,XX, error bars represent SEM. * represent p values calculated from unpaired t test conducted between each experimental and it’s corresponding GFPC

**Supplementary Table 1.**
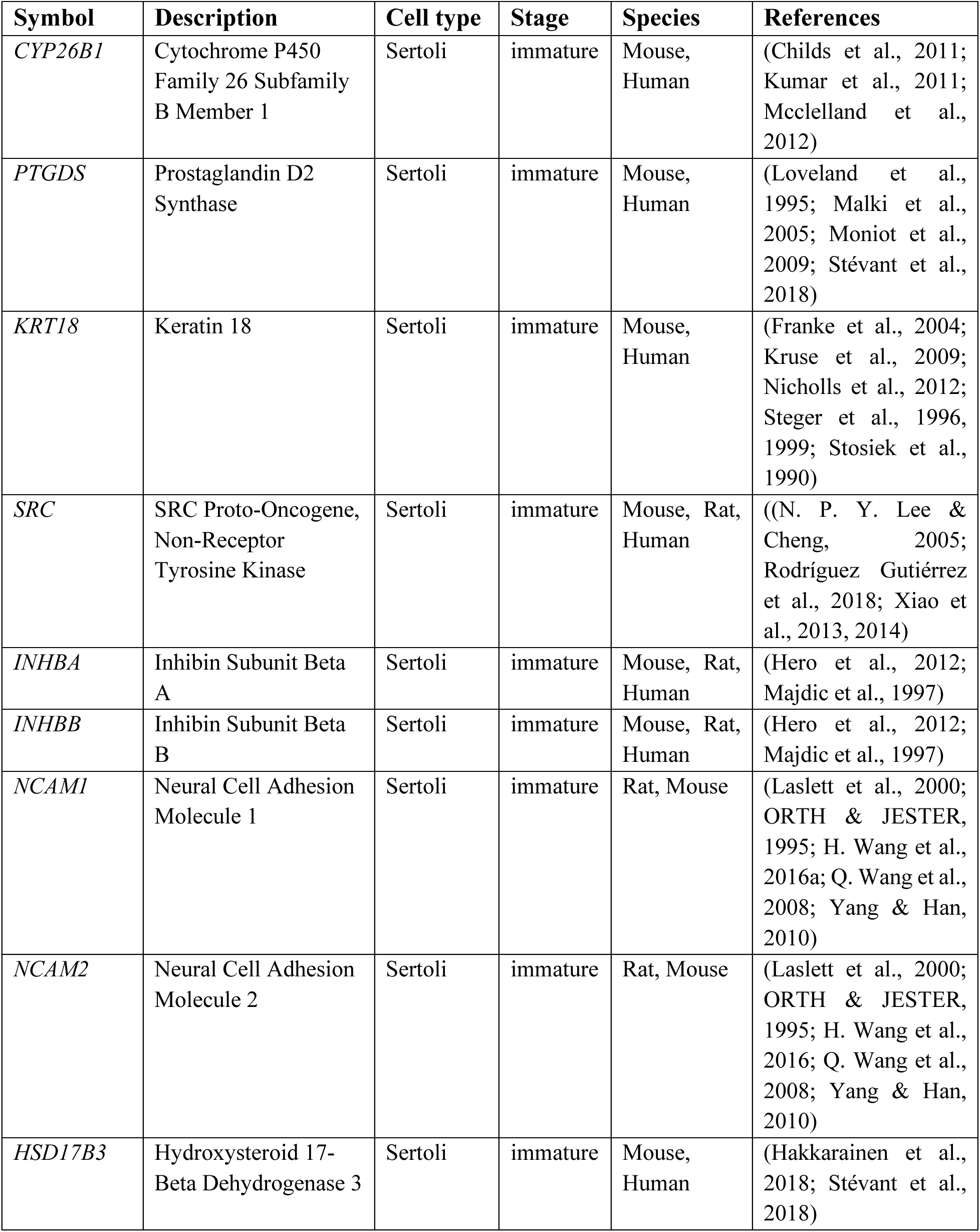
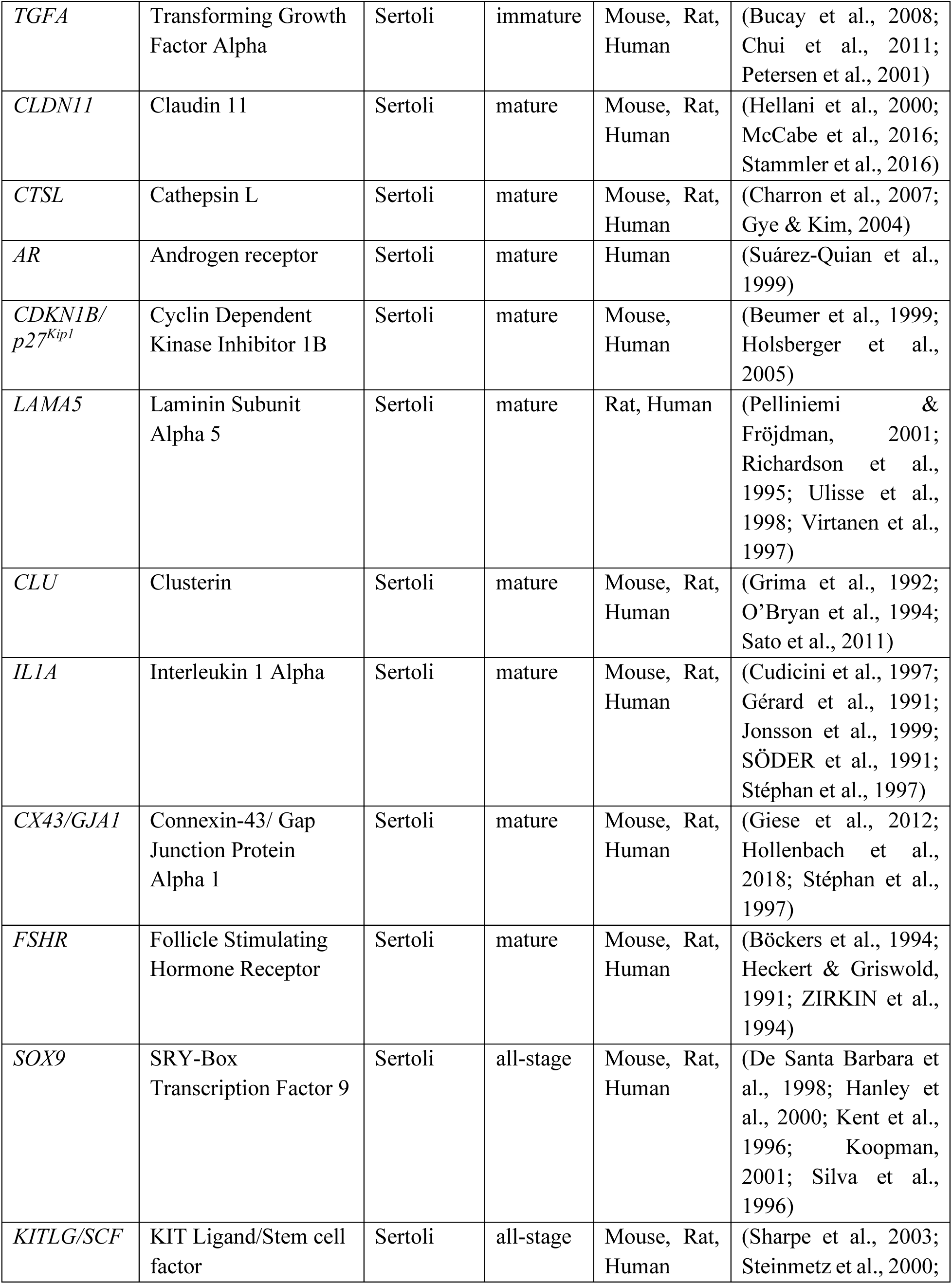
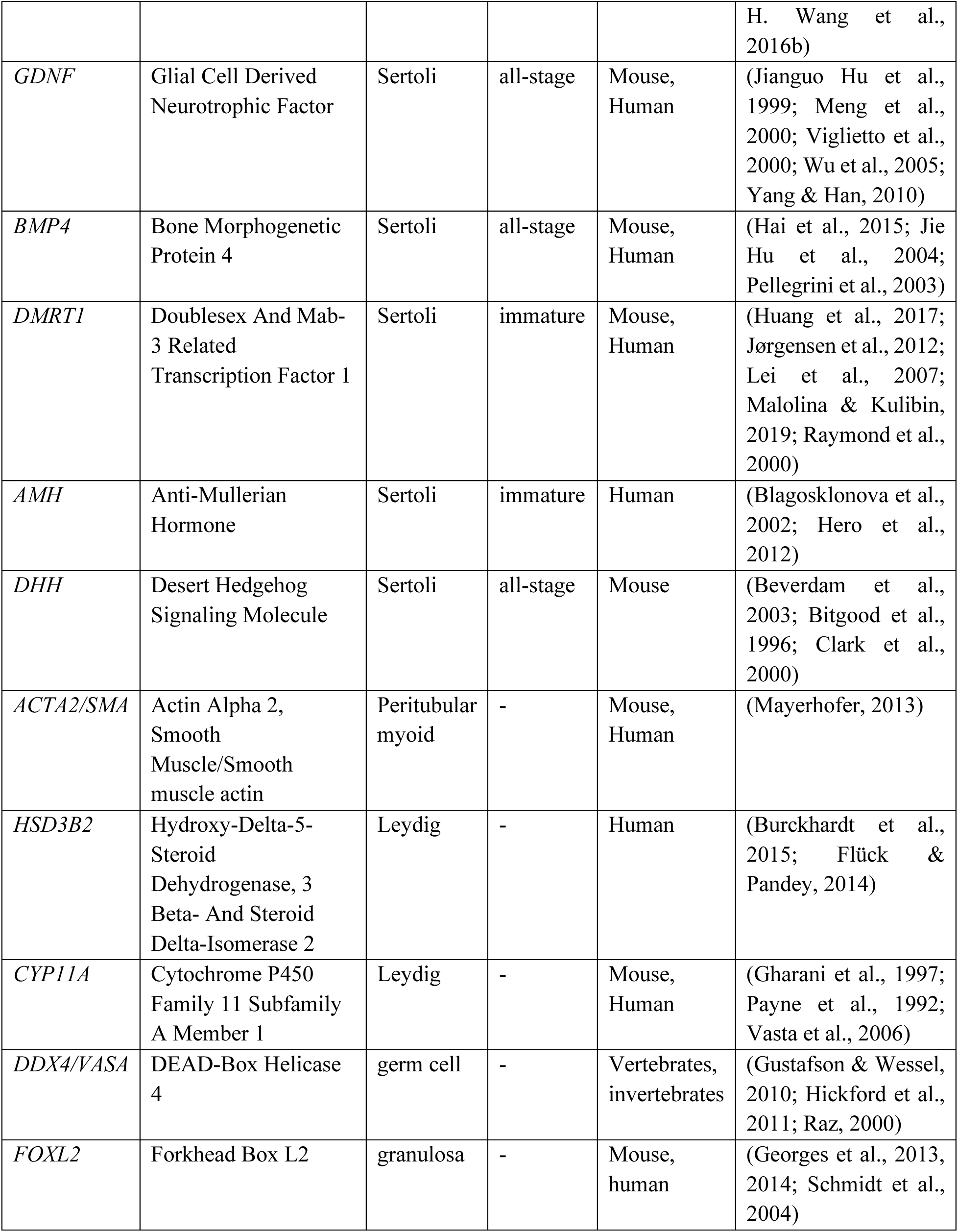
List of marker genes

